# Spatio-temporal summation of perimetric stimuli in healthy observers

**DOI:** 10.1101/2022.08.10.503485

**Authors:** Giovanni Montesano, Pádraig Mulholland, David F. Garway-Heath, Josephine Evans, Giovanni Ometto, David P. Crabb

## Abstract

Spatial summation of perimetric stimuli has been used to derive conclusions about the spatial extent of retinal-cortical convergence, mostly from the size of the critical area of summation (Ricco’s area, RA) and critical number of Retinal Ganglion Cells (RGCs). However, spatial summation is known to change dynamically with stimulus duration. Conversely, temporal summation and critical duration also vary with stimulus size. Such an important and often neglected spatio-temporal interaction has important implications for modelling perimetric sensitivity in healthy observers and for formulating hypotheses for changes measured in disease. In this work, we performed experiments on visually heathy observers confirming the interaction of stimulus size and duration in determining summation responses in photopic conditions. We then propose a simplified computational model that captures these aspects of perimetric sensitivity by modelling the *total retinal input*, the combined effect of stimulus size, duration and retinal cones-to-RGC ratio. We additionally show that, in the macula, the enlargement of RA with eccentricity might not correspond to a constant critical number of RGCs, as often reported, but to a constant critical *total retinal input*. We finally compare our results with previous literature and show possible implications for modelling disease, especially glaucoma.

## Introduction

Measuring how contrast sensitivity varies according to different stimulus sizes and durations has proven invaluable for investigating the psychophysical and physiological basis of transient stimulus detection ^1–5^ and how the underlying physiology is altered by disease^6–10^. In fact, change in sensitivity with increasing stimulus size (*spatial summation*) and duration (*temporal summation*) has been shown to be altered following retinal ganglion cell (RGC) loss from glaucoma^6–9, 11^. Both spatial and temporal summation are characterised by a biphasic response, with a steeper reciprocal relationship between stimulus area/duration and contrast at threshold for smaller/shorter stimuli (total summation) and a shallower change for larger/longer stimuli (partial summation). The response is often characterised in terms of the point of transition between these two phases (*critical size/duration*)^12^. The physiological basis of spatial and temporal summation has been extensively studied. Although models solely based on RGCs exist^13^, spatial summation has been linked to cortical magnification and to the convergence of RGCs onto cells of the visual cortex^10^. This phenomenon is often referred to as *cortical pooling* and it is the favoured model for explaining spatial summation^1, 10^. Cortical pooling can be modelled through linear combination of filter elements tuned to different spatial frequencies^1^.

One aspect that has been explored to a lesser extent is the interaction between stimulus size and duration and its effect on sensitivity (*spatio-temporal summation*). Models exist to describe temporal summation in isolation^14–16^. Many of these authors acknowledge the effect of stimulus configuration^14, 15^ and adaptation state^17^ on critical duration. Direct experimental evidence of the interaction between size and duration for simple circular stimuli^11, 18, 19^ suggests a combined integration of the total input by the visual system. Some attempts have been made to describe such an interaction, mainly in the field of motion detection^20, 21^, but this phenomenon has been little explored for perimetry^19^. Another aspect that has been overlooked is the effect of retinal convergence. One common assumption is that spatial summation at different eccentricities can be exclusively explained by the change in density of RGCs^10^. However, similarly to cortical convergence, individual RGCs’ might carry a different weight in terms of retinal input at different eccentricities because they receive input from a different number of photoreceptors (larger in the periphery), with significant changes in the composition and density of their mosaic

Understanding these aspects is essential for many clinical applications of psychophysics. White-on-white perimetry is one of the most performed tests in clinical practice to diagnose and monitor the progression of a variety of diseases. In its most common implementation, the test is a ‘yes/no’ task in which an observer is asked to press a button every time a stimulus is perceived. The response needs to be provided within a set time window following stimulus onset, with no response indicating that the stimulus was not seen. The stimulus is projected on a bowl with a uniform white background and usually consists of a circular target with sharp edges and 0.43 degrees in diameter (size III according to Goldmann^22^) and a duration between 100 and 200 ms. The intensity of the target is varied to estimate the 50% seen contrast threshold, using a variety of strategies. The target is presented at various locations around the fixation target, according to a set of pre-determined testing grids, so that the 50% threshold can be estimated at each of these locations. This produces a sensitivity map that can be used to identify and monitor visual field defects. The objective of our work was to collect experimental data to build and validate a spatio-temporal summation model, able to capture the combined effect of retinal convergence, stimulus size and stimulus duration for perimetric stimuli.

## Methods

### Participants

Ten visually healthy participants between 18 and 40 years of age were recruited on a voluntary basis at City, University of London, London, United Kingdom. All participants gave their written informed consent. The study was approved by the local Ethics board (Optometry Proportionate Review Committee, approval number ETH2021-1728) and adhered to the tenets of the declaration of Helsinki. All participants underwent an ophthalmic assessment by an ophthalmologist (GM), which included objective refraction and measurement of the intraocular pressure (IOP) with a non-contact tonometer and auto-refractor (TRK-1P, Topcon, Tokyo, Japan), best corrected visual acuity (BCVA) with Snellen charts, slit lamp assessment of the anterior segment and indirect fundoscopy. Reasons for exclusion were any abnormality of the retina or of the optic nerve head (ONH), IOP > 21 mmHg and a BCVA < 6/6 in the test eye. If both eyes were eligible, the one with the smallest refractive error was selected.

### Psychophysical experimental procedure

#### Testing apparatus

All experiments we carried out on an Octopus 900 bowl perimeter (Haag Streit AG, Koeniz, Switzerland) controlled through the Open Perimetry Interface^23^. The bowl is 30 cm in radius. The perimeter is equipped with a chinrest and an infrared camera to monitor eye position and pupil size. Chinrest position was adjusted by the operator as required, to maintain good centration of the pupil. A central target (four small dots in a diamond arrangement) encouraged fixation and avoided interference with centrally presented stimuli. A near-vision lens addition of approximately +2.50 D was used to reduce strain from accommodation, refined with subjective assessment of optimal visibility by the subject. Lenses were placed on an adjustable lens holder in-built to the instrument. The background illumination was 10 cd/m^2^. Calibration was performed in a dark room before every experiment through an automated procedure implemented by the manufacturer. As it is convention in perimetry, the intensity of the stimulus in dB is expressed as attenuation of the maximum possible stimulus intensity (3185 cd/m^2^), so that higher contrast equates to lower dB values. This quantity can be converted to Weber contrast (*W_c_*) using Equation (1). However, for simplicity in our calculations, we report the values as Differential Light Sensitivity (DLS), which is simply the sensitivity value in dB/10.

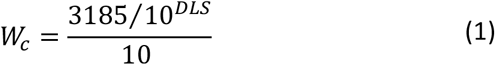

#### Spatio-temporal summation

In the first experiment, we estimated contrast sensitivity at twelve locations in the central visual field (VF) with different stimulus sizes and durations for one test eye of all ten participants. The locations’ coordinates ({X; Y}) in visual degrees from fixation were: {±7; ±7}; {±4, ±4}; {±1, ±1}. Stimuli were round achromatic targets with five different diameters (Goldmann sizes, G): 0.10 (G-I); 0.21 (G-II); 0.43 (G-III); 0.86 (G-IV); 1.72 deg. (G-V). All locations were tested with all stimulus sizes. The locations at {±7; ±7} were additionally tested with five different stimulus durations (for all stimulus sizes): 15 ms; 30 ms; 55 ms; 105 ms; 200 ms. Four combinations (G-I/15 ms; G-I/200 ms; G-V/15 ms; G-V/200 ms) were tested twice so that more robust estimates of their threshold were available for the measurement of the frequency of seeing (FOS) curves (see next section).

The threshold was determined with a yes/no task. The observer was asked to press a button every time a stimulus was perceived. We assumed that no response within a predetermined time widow (1500 ms) corresponded to “not seen”. The threshold was estimated through a Bayesian strategy, the Zippy Estimation through Sequential Testing (ZEST)^24^, as implemented on the OPI. For our test, the strategy was set to have a uniform prior distribution between 0 and 50 dB (the range of the instrument). The likelihood function was a Gaussian cumulative distribution function (CDF) with a standard deviation (SD) of 1 dB and a guess/lapse rate of 3%. The prior distribution was updated at each response to generate a posterior distribution. The posterior distribution was used as the prior distribution for next step in the strategy. The stimulus was chosen as the mean of the prior distribution at each step, rounded to the closest integer dB value. This has been shown to provide unbiased estimates of the 50% detection threshold for a yes/no task^24^. The determination of each threshold terminated when the posterior distribution reached a standard deviation < 1.5 dB (dynamic termination criterion).

Each combination of stimulus size and duration at each location was treated as a separate independent “thread” by the strategy (140 in total). The threads were randomly subdivided into four blocks, to allow for breaks within the test. Each block of testing lasted for approximately 15 minutes (~350 presentations). Individual presentations within each block were fully randomised. A block was completed when all the 35 threads assigned to it reached the termination criterion. A pause between individual presentations was also introduced, calculated as (1000 ms – response time, minimum 200 ms) plus an additional pause, randomly sampled from a uniform distribution between 0 and 100 ms. All responses occurring within the pause or less than 180 ms after stimulus onset stimulus ^25^ were considered as false responses and discarded.

#### Frequency of seeing curves

For a subset of five participants, frequency of seeing (FOS) curves were determined for four stimulus combinations (G-I/15 ms; G-I/200 ms; G-V/15 ms; G-V/200 ms) at coordinates {±7; ±7} degrees (four locations) using a Method of Constant Stimuli (MOCS) procedure. Following others^8^, we used a two-stage approach. First, we obtained a coarse estimate of the FOS curve through a multidimensional Bayesian strategy, QUEST+^26^. Such a strategy is similar in principle to ZEST but uses entropy to determine the next presentation and allows for multiple parameters to be estimated. In our procedure, the FOS curve was parametrised as the CDF of a Gaussian distribution, with a fixed guess/lapse rate of 3%. The mean and SD (which model the 50% threshold and the slope of the FOS curve respectively) were simultaneously fitted as free parameters. The test was terminated when the entropy of the combined posterior distribution was ≤ 4.5. For the purpose of this preliminary step, the four spatial locations were considered as interchangeable. Therefore, only four FOS curves were determined, one for each stimulus combination. The prior distribution for the mean was itself a Gaussian distribution with a SD of 4 dB, centred on the average of the sensitivity estimates obtained from the ZEST procedure for the tested locations (8 estimates for each stimulus combination, i.e. 4 locations each tested twice) and limited over a domain of ±5 dB around its mean. The prior distribution for the SD of the FOS curve was a uniform between 1 and 10 dB, with steps of 0.5 dB.

The estimated SD for the Gaussian FOS curves were used to determine the contrast levels to be tested for each stimulus combinations in the actual MOCS. We tested seven steps for each location and each condition. The steps were placed at the following quantiles of the Gaussian FOS (neglecting lapse/guess rate): {0.0001, 0.1, 0.3, 0.5, 0.7, 0.9, 0.9999}. We however ensured that all the steps were at least 1 dB apart (the minimum interval allowed by the device) and that the two most extreme contrast levels were at least 10 dB above and below the estimated 50% threshold. The 50% threshold was calculated as the average of the two test results obtained from the ZEST strategy for each location. Each contrast level was presented 25 times and each spatial location was tested fully and independently, for a total of 2800 presentations. A break of at least 10 minutes was introduced every 350 presentations and the whole test was split into two sessions performed on two separate days. The individual presentations were fully randomised across test locations, stimulus area/duration combinations and contrast levels. Pauses between presentations and false responses were determined as described above for the main experiment.

MOCS data were fitted using a Bayesian hierarchical model, similarly to Prins^27^. The results of the test performed on each subject were fitted independently. The psychometric function was modelled with the CDF of a Gaussian function (Φ), where the mean (μ), SD (σ), lapse rate (λ) and guess rate (y) were free parameters (see **Equation 2**). Mean (μ) and σ were hierarchical parameters that varied for each of the four tested locations. Information was however propagated across different locations to improve the robustness of the fit of the parameters for each testing condition. Lapses and guesses were instead modelled as global parameters for the whole test. Details of the implementation of the Bayesian model are reported in the **Appendix**.

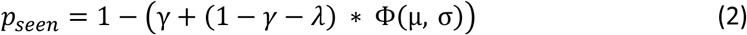

### Imaging

Retinal imaging was performed using a Spectralis Spectral Domain Optical Coherence Tomography (SD-OCT, Heidelberg Engineering, Heidelberg, Germany) scanner. Dense macular volume scans spanning the central 25 x 30 visual degrees (121 vertical B-scans, 9 averaged scans) were segmented and exported as RAW files using the Heidelberg Eye Explorer (HEYEX, Heidelberg Engineering, Heidelberg, Germany). Retinal ganglion cell layer (RGCL) thickness maps were built from segmentation data and converted to customised estimates of local RGC counts by combining thickness data with histology data provided by Curcio and Allen^28^, using previously published methodology^29, 30^. Local customised RGC density was calculated for each location tested in the psychophysical procedure by accounting for RGC displacement^30, 31^, using methodology detailed elsewhere^30^.

### Modelling of perimetric sensitivity

One of the objectives of this study was to provide a model that was simple, but sufficient to describe the change in sensitivity observed with different combinations of sizes and durations for perimetric stimuli. Our working hypothesis, derived from previous work^11, 18, 19, 32^, was that the combined effect of these two parameters, at any given location, could be described by taking the product of stimulus area and stimulus duration. We called this product the *spatio-temporal input*. We integrated the spatio-temporal input into a computational model of the response of RGC mosaics, partially based on the work by Pan et al.^1^ and Bradley et al.^33^. The key novel aspect of our modelling was that the linear response from the RGC mosaic was pooled and integrated over time so that changes in duration and size of the stimulus would both simultaneously affect the temporal and spatial response of the system. We further modelled the retina as a two-stage mosaic, where the response from individual photoreceptors active in photopic adaptation conditions (cones) was integrated by the RGC mosaic, to explore the effect of retinal convergence in the central visual field. The density of the two mosaics was varied to reproduce the effect of eccentricity. We refer to the combined effect of the spatio-temporal input and changes in retinal structure (i.e. density of the photoreceptor and RGC mosaics) as *total retinal input*. The model was implemented in Matlab (The MathWorks, Natick, USA) and is described in detail below.

#### Hexagonal mosaics

Following Swanson et al.^7^, we modelled multiple detectors organised in a regular hexagonal lattice. This organisation is reflective of many naturally occurring cell mosaics as it represents the most efficient packing scheme for objects with circular/spherical geometries^34^. For our purposes, we simplified the retina as being composed of two stacked mosaics, the photoreceptor mosaic and the RGC mosaic. Being interested in the results of experiments performed in photopic conditions (background illumination = 10 cd/m^2^), we only modelled the cone mosaic. In this retinal model, individual RGCs pool the response from the photoreceptors according to their Receptive Fields (RFs). To improve the efficiency of computation, each hexagonal lattice was rearranged in a regular lattice with anisotropic spacing (See **Appendix Figure 1**). This simplifies the pooling operation, which can be computed via simple convolution of the regularised lattice with the RGC-RF filter (see next section), also rearranged according on the same regular lattice. The response of the photoreceptor mosaic was simply computed by multiplying the mosaic by the stimulus. In its simplest form, this is equivalent to assigning a value of 1 to all the photoreceptors that fall within the stimulus area, leaving the others to 0. However, in its final implementation, this was modified to include the effect of optical blur (see later). Only the midget OFF RGC mosaic was used for the calculations (mOFF-RGC), assuming that the ON and OFF mosaic operate on parallel redundant channels for the detection of simple round stimuli.

#### RGC receptive field

The spatial filters for the RGC-RF were modelled with a Difference of Gaussian (DoG, **Figure 1 A**), using the median parameters estimated by Croner and Kaplan^35^ from electrophysiology on macaque’s retina. In their work, they showed that, although the scaling factors for the relative width and height of the inhibitory and excitatory Gaussian components of the filter changed with eccentricity, their ratios remained approximately constant. In this model, the surround inhibitory component has peak sensitivity *K_s_* = 0.01\**K_c_*, where *K_c_* is the peak sensitivity of the excitatory centre. The standard deviation (SD) of the surround was 6.7 times larger than the SD for the centre (average reported by Croner and Kaplan^35^). The SD for the centre was scaled so that the *radius* of the centre component was equal to the inter-cell spacing of the mosaic (defined by its density). The radius was defined by Croner and Kaplan as the distance from the centre at which the excitatory Gaussian component has value *K_c_/e*. The corresponding SD was approximated as SD = Cell spacing/1.414. Note that, while the centre-surround proportions are based on Croner and Kaplan^35^, the actual extent of the RGC-RFs in our model depends only on the inter-cell spacing of the RGC mosaic.

**Figure 1.**
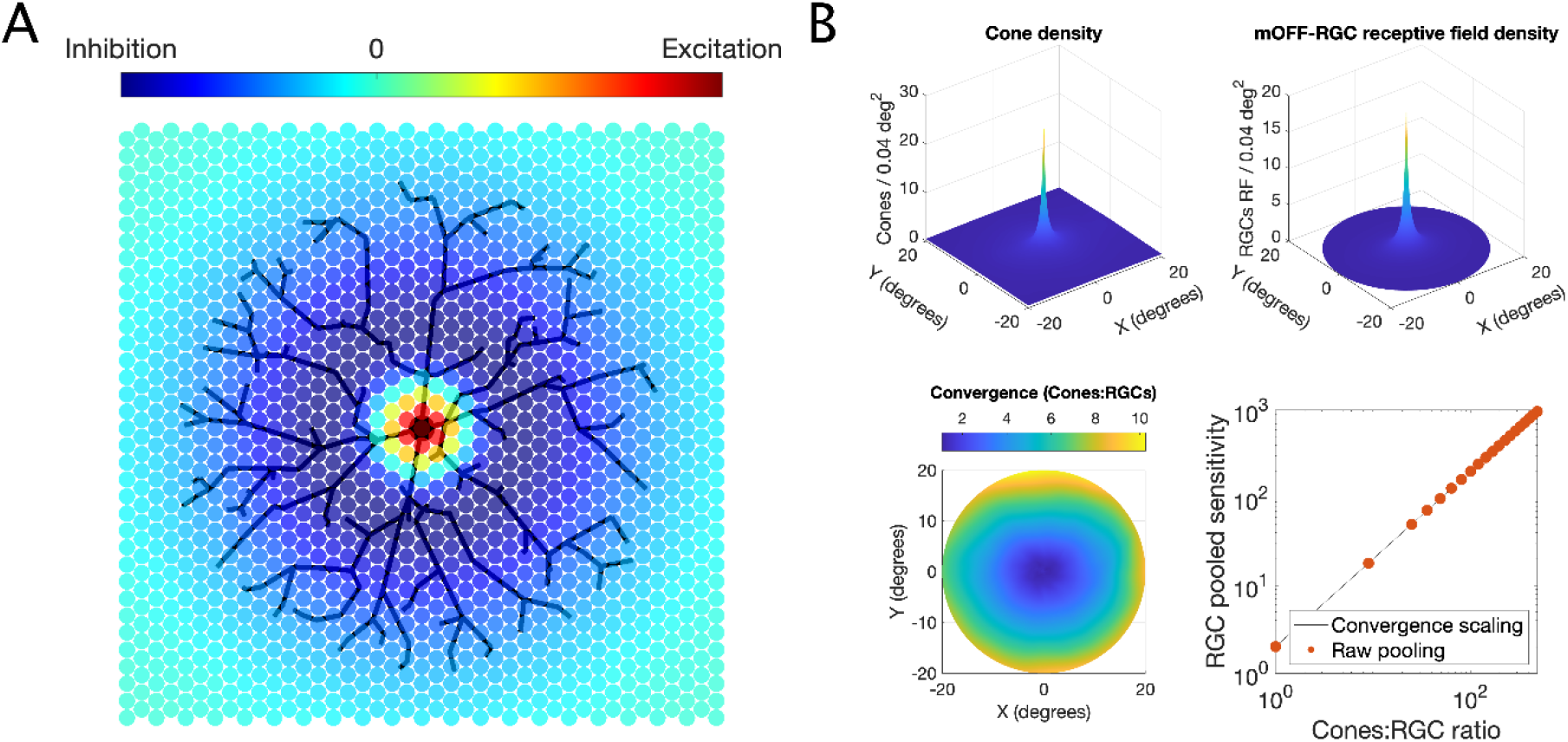
**A)** Schematic example of how a Retinal Ganglion Cell (RGC) samples the input from the photoreceptor mosaic, according to its Difference-of-Gaussian receptive field (RF). The strength of inhibitory surround has been exaggerated here for clarity. **B)** Estimated density for Cones (top left) and RGC-RF (top right) and a map of Cones:RGCs convergence (bottom left). The bottom-right panel shows a comparison between the predicted (unscaled) sensitivity for the numerical calculations from the mosaic with discrete changes in convergence (dots) and continuous factor scaling (line).

#### Cone-RGC convergence

The number of cones that converge onto a RGC is known to increase with eccentricity^28, 31, 36^. In our model, this corresponds to an increasing number of photoreceptors pooled by the RGC-RF per unit area. This can be achieved by increasing the density of the cone photoreceptor mosaic, also provided by Curcio and Allen^37^. The convergence rate can be calculated by taking the ratio of the density of cones over the density of mOFF-RGCs (**Figure 1 B**). As expected, the smallest value for convergence was very close to 1 (1.08), near the fovea. Because of how the hexagonal matrix has been re-arranged for calculations (fig.1), the intercell spacing for the RGC mosaic needs to be an exact multiple of that of the cone mosaic. This limits the possible Cones:RGCs ratios that can be calculated. However, changing the convergence ratio is equivalent to simply multiplying the response of the RGC obtained with a 1:1 convergence ratio by a scaling factor. This is easily demonstrated by the graph in **Figure 1 B**. This method was therefore chosen to account for the change in convergence across the VF in a smooth fashion.

#### Modelling of optical factors

The effect of natural optics was modelled using the formula for the average Modulation Transfer Function (MTF) of the human eye proposed by Watson^38^. In this formula, the square-root of the diffraction limited (DL) MTF, which depends only on the pupil size, is multiplied by a Lorentzian function whose parameters are fitted so that the product would approximate the average human MTF. A multiplicative correction factor, that depends on age and eye pigmentation, is then additionally applied to the MTF to account for light scattering. **Figure 2** reports examples of the effect of optical blur on different stimulus sizes for different pupil apertures using the MTF (without accounting for scattering)^38^. The calculations are performed by multiplying the two-dimensional Fourier transform of the stimulus by the MTF and then back-transforming in the spatial domain. The blurred stimulus can then be sampled with the photoreceptor mosaic. For each subject, we used the average pupil size recorded by the Octopus perimeter during the test to model the results of our experiments.

**Figure 2.**
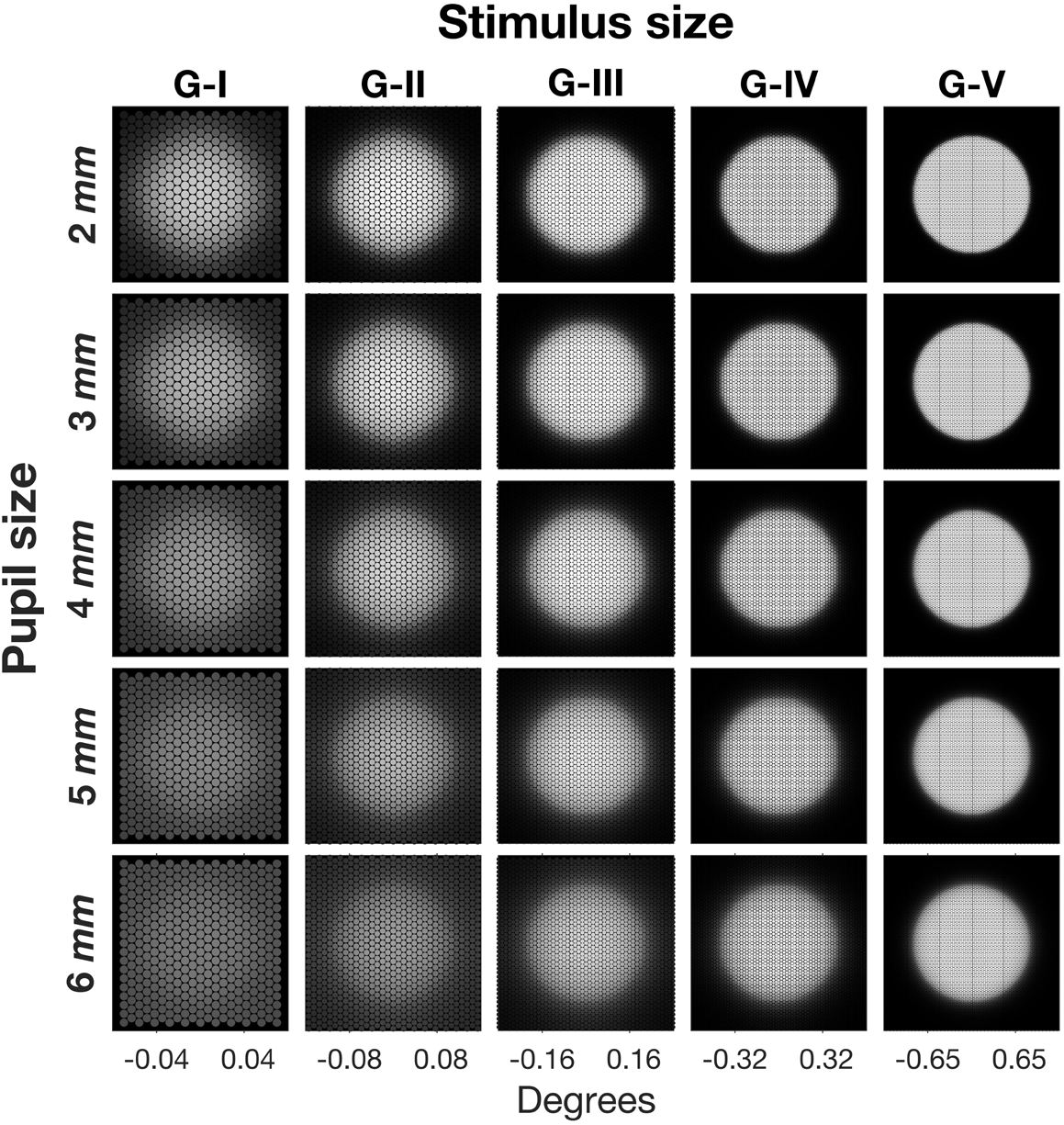
Effect of optical blur for different pupil sizes. The images represent the projection of the blurred stimulus on the photoreceptor mosaic.

#### Proposed spatio-temporal model

One desired property of our proposed model was that the size and duration of the stimulus interacted so that longer stimuli would decrease Ricco’s area (upper limit of complete spatial summation) and larger stimuli would shorten the critical duration (upper limit of complete temporal summation). One solution to achieve this is to use a pooling operation that integrates the spatial input over time. The integration, however, must not solely take into account the duration of the stimulus, but also the amount of RGCs stimulated (i.e., the amount of spatial input). In other words, the temporal integration is to be performed by a cortical pooler on the total spatial input rather than by individual detectors prior to pooling. The simplest model, with the smallest number of parameters, is a capacitor (Equation 5), which is convolved with the temporal profile of the stimulus and then integrated over time according to Equation (6) to obtain the responsed

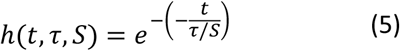

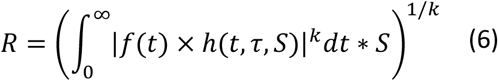

where *τ* is the integration constant, *k* is the summation exponent (4 in this study)^1, 7, 39–42^ and *S* is the total spatial input defined as

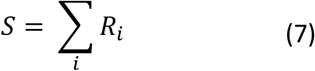

where *R_i_* is the response of an individual ganglion cell to the stimulus. Note the contribution of individual RGCs (*R_i_*) can change both because of the location of the RGC with respect to the stimulus (edge as opposed to centre) and the effect of retinal convergence (RGCs in the periphery will have a bigger contribution when fully stimulated because of their larger pooling from the photoreceptors). The temporal profile of the stimulus is represented by *f*(*t*), which is a step function with value 1 when the stimulus is on and 0 otherwise. As previously mentioned, the combined effect of stimulus size, stimulus duration, RGC density and retinal convergence defines the *total retinal input*. Much like other temporal filters, this operation can also be implemented through temporal convolution. Note that such an approach to spatio-temporal summation is very similar to what was described in Frederiksen et al.^21^ and Anderson and Burr^20^ for motion detection. Since only the mOFF-RGC mosaic was considered for our calculations, the RGCs that were assigned a negative input were considered as inhibited by the stimulus. Their negative contribution to the sum can be interpreted as an inhibition of their background activity. Obviously, such a simple approach would not account for other filter choices with a strong biphasic response, where a simple summation would always result in a zero net sum. From the examples in **Figure 3**, we can see that this pooler has the desired properties when the response is computed for different stimulus sizes and durations, i.e. a shorter duration determines a larger critical area and vice-versa. One additional convenient property of this pooler is that the critical size and duration depend on the integration constant τ. The integration constant τ is therefore the scaling factor of the pooler and can be used to test the hypothesis of constant input integration across the VF. If the hypothesis of constant integration response for the same amount of total retinal input is correct, we do not expect important changes in the integration constant across different testing conditions and eccentricities. An alternative approach would be to model individual RGCs (or higher order visual detectors) as separate spatio-temporal integrators and to pool their response by vector summation^1, 39^. Such an approach has the advantage of allowing the modelling of the response from specific classes of RGCs and produces sensible spatial and temporal summation responses. However, it fails to reproduce the interaction between spatial and temporal input that would be expected. For example, Ricco’s areas in spatial summation curves would be unaffected by changes in stimulus duration. This is in contrast with evidence from the literature^11, 18, 19, 32^. It is worth noting that the current model could be extended to include the temporal response of individual classes of RGCs prior to pooling. However, this would increase the number of tuneable parameters and would be beyond the objectives of the current study and what could be determined with our experiments.

**Figure 3.**
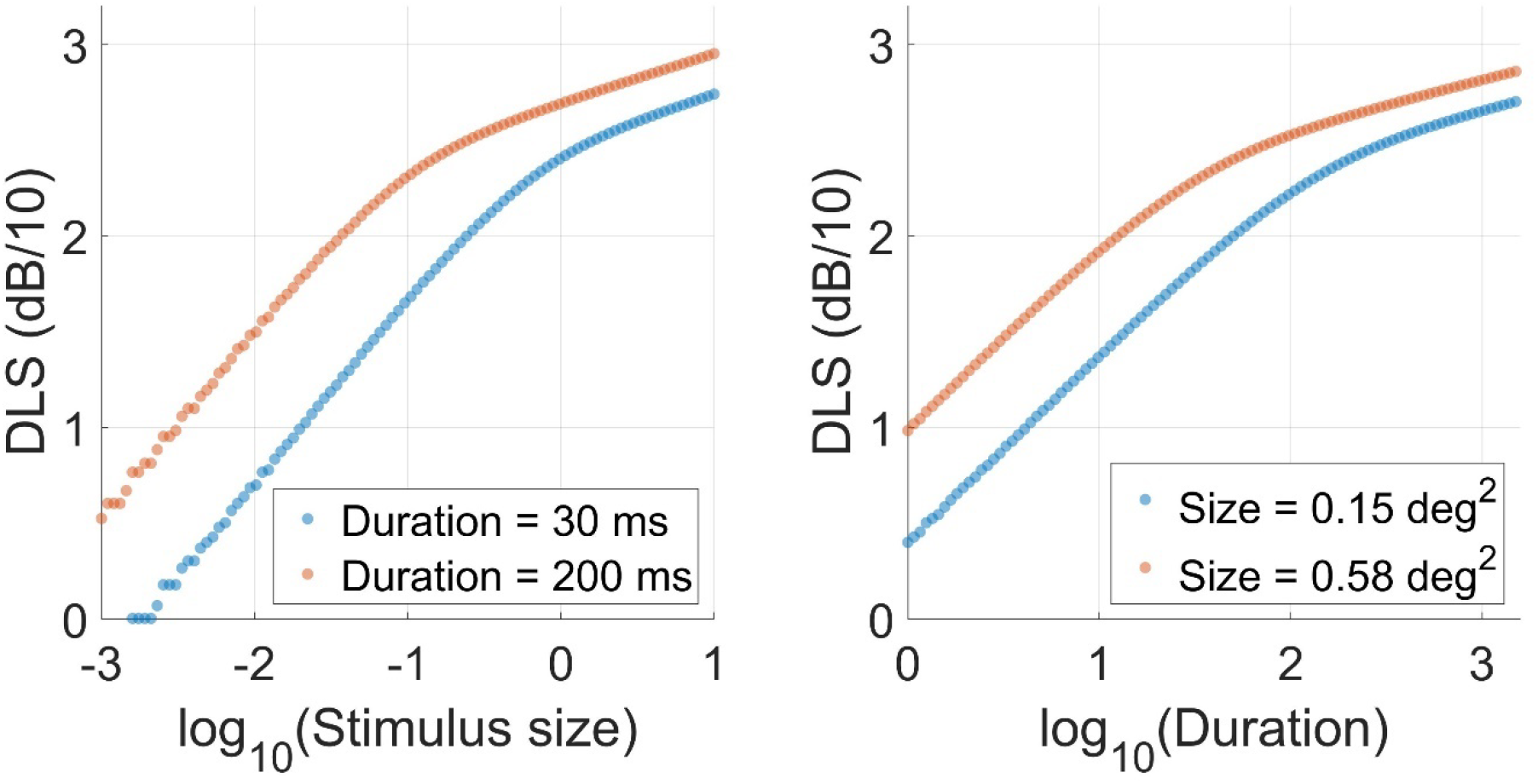
Example of the interaction of stimulus size and duration in the proposed model. Changing the stimulus duration translates the spatial summation curve along the horizontal axis (left panel). The same is true for the temporal summation curve when changing the stimulus area (right panel).

#### Fitting procedure

The model described by Equation (6) was fitted to the data using an iterative algorithm (Nelder-Mead Simplex Method, *fminsearch* function in Matlab^43^) to minimise the Root Mean Squared Error (RMSE). The summation exponent was set to *k* = 4^1, 7, 39–42^ and the RGC mosaic density was varied according to the eccentricity following the model by Drasdo et al.^30, 31^. These estimates were corrected with individual imaging data obtained from the OCT scans, as previously reported^29, 30^. The model was fitted by tuning the parameter τ, which represents the integration constant of the spatio-temporal input. An additional parameter (additive in log-scale) allowed translation along the vertical axis (log-DLS, Offset term).

#### Calculation of critical size

The transition from total to partial summation is smooth for the curves generated by our model. The response curve is fully characterised by the integration constant *τ* and the amount of retinal input. The calculation of the critical (Ricco’s) area is therefore dependent on an arbitrary threshold and is only performed for comparison with previous literature. For our calculations, the transition point was the retinal input at which the slope of the summation curve is 0.5 (Piper’s law). Note that the retinal input scales perfectly with stimulus size for our chosen implementation of the model, but non linearities are introduced if taking the sum of the module in Equation (7). For consistency with our supplementary analyses (see later), the conversion between stimulus area and retinal input for each mosaic was calculated numerically and locally approximated with a linear function in log_10_ - log_10_ scale. The parameters for the curves were fitted accounting for the optical blur (based on each participant’s average pupil size and iris pigmentation). Densely sampled curves were numerically calculated using these parameters to estimate Ricco’s area. These curves were calculated without the effect of optical blur. This simulates removing the estimated effect of optics on the size of Ricco’s area. Note that accounting for convergence in the fitting process will not change Ricco’s area, as parameters are optimised to fit the same data.

#### Statistical analysis

Statistical comparisons were performed using linear mixed models to account for correlations between observations from the same subject. When data from multiple locations were used, individual locations were used as a nested random factor within the subject. When multiple comparisons were compared, the p-values were corrected using a Bonferroni-Holm correction. All calculations were performed in R (R Foundation for Statistical Computing, Vienna, Austria) using the lme4 package^44^. All comparisons were performed on log10-transformed values of Ricco’s area, integration constant and number of mOFF-RGCs, unless otherwise specified. Eccentricity was treated as a discrete factor.

### Results

#### Average response

In this section, we show plots of the average DLS for different experimental conditions to give an intuitive representation of the phenomena under investigation. Characteristics of each eye in the sample are reported in **Table 1. Figure 4A** reports the average DLS for the spatial summation experiment at different eccentricities. As expected, the summation curves are separated by a horizontal shift, owing to the effect of the changes in the retinal mosaic. Interestingly, simply transforming the stimulus area into the corresponding estimated number of RGC-RFs underlying the stimulus did not fully account for the effect of eccentricity. Most of the effect was instead removed by considering the product of stimulus area, RGC-RF density and Cones:RGC convergence ratio. We evaluated this by comparing the results of a simple 2^nd^ degree polynomial fit of the DLS using either the log_10_(stimulus area), the raw log_10_(number of RGCs) or the convergence weighted log_10_(number of RGCs) as predictors in a mixed effect model. The unexplained residual variance (including random effects) was 1.93 dB^2^ for the log_10_(stimulus area), 1.84 dB^2^ for the unweighted log_10_(number of RGCs) (4.4% reduction) and 1.77 dB^2^ for the convergence weighted log_10_(number of RGCs) (8.2% reduction).

**Figure 4.**
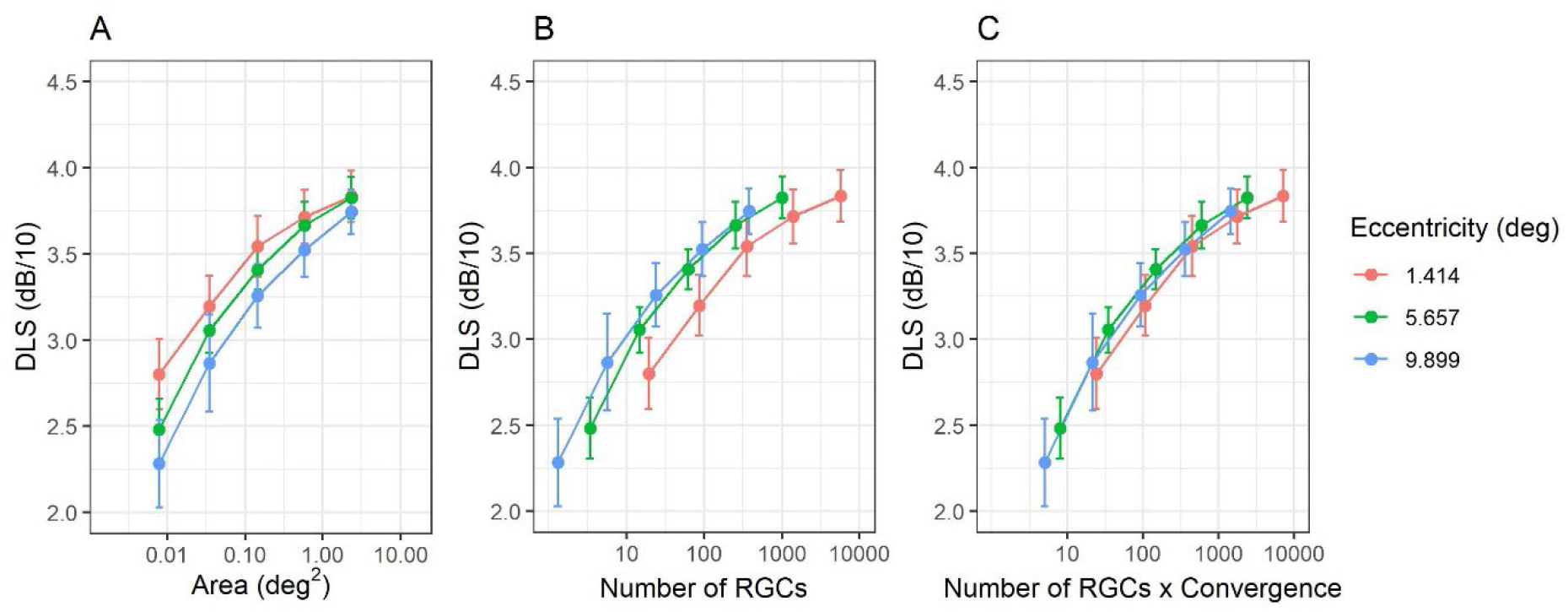
Average (dots) and standard deviation (error bars) for Differential Light Sensitivity (DLS) for the three tested eccentricities at different stimulus sizes **(A)**, the corresponding mOFF-RGC-RF count underlying the stimuli **(B)** and the corresponding mOFF-RGC-RF count underlying the stimuli weighted by convergence **(C)**. RGC = Retinal Ganglion Cell (average across subjects at each stimulus size in these graphs).

**Table 1.** Characteristic of each eye in the sample. All subjects had their sensitivity tested with the ZEST strategy for all the duration and size combinations for all tested locations. Psychometric functions were estimated for subjects from 1 to 5 using the method of constant stimuli. D = Diopter; BCVA = Best Corrected Visual Acuity; logMAR = log-Minimum Angle of Resolution; IOP = Intraocular Pressure; GCL = macular Ganglion Cell Layer; RNFL = peripapillary Retinal Nerve Fibre Layer. Average macular and GLC thickness were measured for the area corresponding to the central 10 degrees.

| Subject ID | Age (years) | Study eye | Sphere (D) | Cylinder (D) | Axis (deg) | BCVA (logMAR) | IOP (mmHg) | Average Macular thickness (μm) | Average GCL thickness (μm) | Average RNFL thickness (μm) |
| --- | --- | --- | --- | --- | --- | --- | --- | --- | --- | --- |
| Subject 1 | 33 | Left | -3.00 | -1.00 | 154 | 0.02 | 16 | 306.4 | 37.9 | 110.9 |
| Subject 2 | 25 | Right | +0.25 | -0.75 | 31 | -0.10 | 14 | 308.1 | 39.7 | 92.2 |
| Subject 3 | 33 | Left | -3.25 | -0.25 | 111 | 0.01 | 18 | 339.5 | 42.5 | 106.2 |
| Subject 4 | 27 | Left | -0.25 | -0.50 | 171 | -0.10 | 14 | 330.0 | 42.2 | 98.9 |
| Subject 5 | 25 | Left | +0.75 | -0.75 | 8 | 0.00 | 15 | 311.3 | 37.4 | 111.9 |
| Subject 6 | 26 | Right | -0.25 |  |  | 0.01 | 11 | 311.3 | 40.9 | 104.6 |
| Subject 7 | 36 | Left | +0.25 | -1.00 | 173 | 0.00 | 19 | 314.7 | 42.7 | 104.6 |
| Subject 8 | 28 | Right | -2.25 | -0.50 | 43 | 0.00 | 15 | 298.6 | 33.7 | 81.5 |
| Subject 9 | 26 | Right | -0.75 | -0.25 | 7 | 0.02 | 16 | 311.1 | 38.0 | 105.6 |
| Subject 10 | 32 | Right | -2.00 |  |  | 0.00 | 15 | 295.4 | 40.8 | 90.3 |

**Figure 5** reports the average DLS for the different testing conditions at locations {±7; ±7} and shows how both spatial and temporal summation curves are affected by changes in stimulus duration and size respectively. However, the values seem to follow a common trend when plotted according to the spatio-temporal input, i.e. the product of stimulus area and duration. We evaluated this by comparing the results of a simple 2^nd^ degree polynomial fit of the DLS using either the log_10_(stimulus area) or the log_10_(spatio-temporal input) as predictors in a mixed effect model. The unexplained residual variance (including random effects) was 11.4 dB^2^ for the log_10_(stimulus area) and 3.7 dB^2^ for the log_10_(spatio-temporal input), a 67.5% reduction.

**Figure 5.**
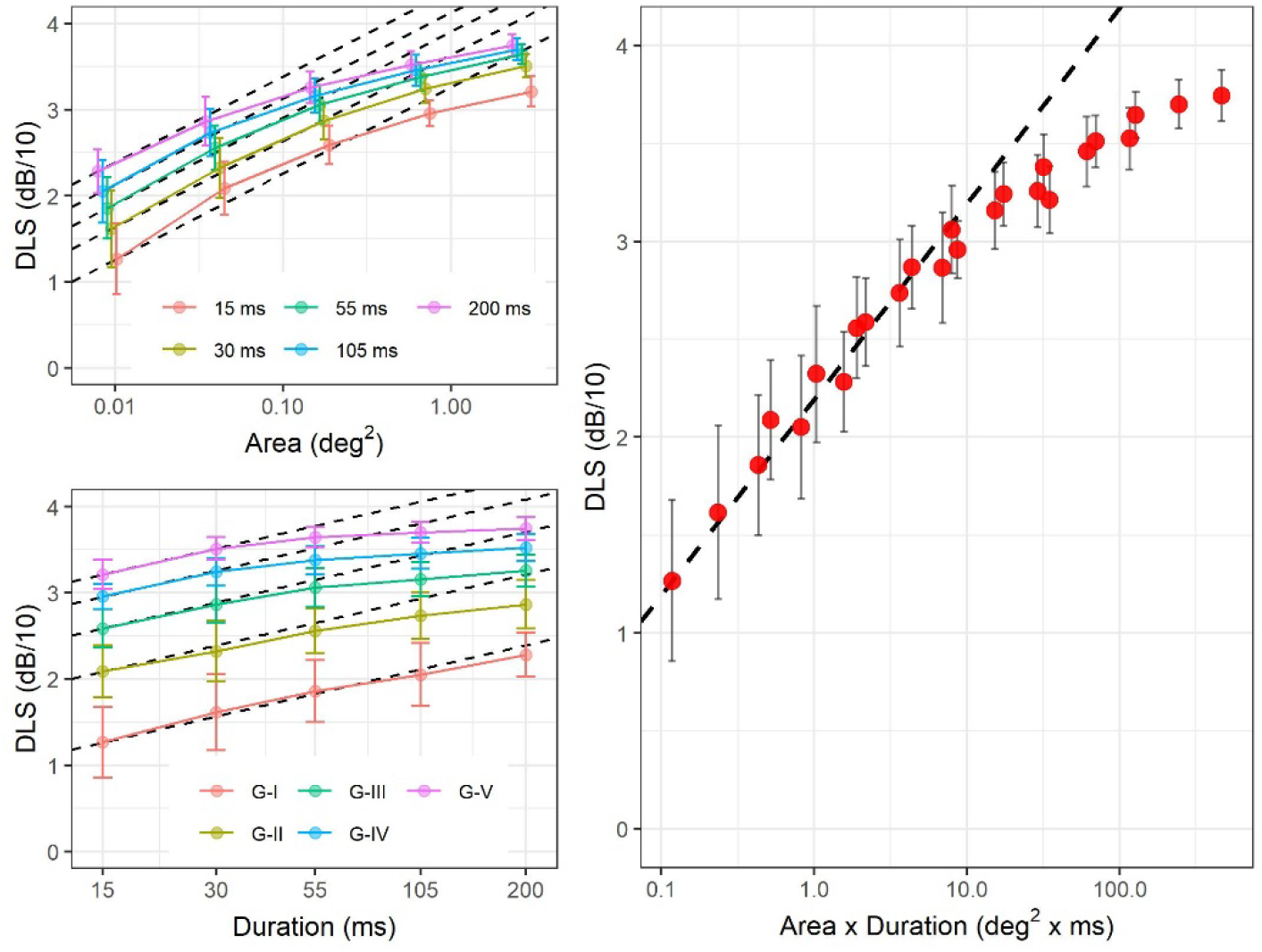
Average (dots) and standard deviation (error bars) for Differential Light Sensitivity (DLS) at the largest eccentricity with different combinations of stimulus sizes and duration. The dashed line indicates total (complete) summation, with intercept equal to the smallest mean value. A small horizontal shift was added to the spatial summation plots to improve visibility.

Taken together, these results and plots support the hypothesis that the main determinant of DLS is the total retinal input to higher visual centres, influenced by both the number of stimulated RGCs, retinal convergence and duration of the stimulus.

#### Results from the spatio-temporal model

##### Spatial summation – effect of eccentricity

The parameters of the model were fitted independently for each location using the data collected with different stimulus sizes and 200 ms stimulus duration (the only duration tested at all eccentricities). **Figure 6** reports the estimated critical size (Ricco’s area) at different eccentricities. The average root mean squared error (RMSE) of the model fits was 0.85 ± 0.39 dB (Mean ± SD). As expected, the estimated Ricco’s area increased towards the periphery (**Figure 7C** and **Table 2**), with no significant differences between the areas calculated with and without accounting for convergence. However, such a change did not correspond to a constant number of mOFF-RGCs being stimulated. Instead, the estimated number of mOFF-RGCs at Ricco’s area was consistently larger towards the fovea (**Figure 6D**). This was mirrored by a change in the integration constant τ with eccentricity. However, this trend in τ was completely eliminated by accounting for the change in Cones:RGCs convergence (**Figure 6 A and Table 2**). This result can alternatively be visualised by multiplying the number of mOFF-RGCs at Ricco’s area by the corresponding convergence factor (**Figure 6D and Table 2**). Note that this is a post-hoc calculation and not an output from the model (accounting of convergence is expected to have an effect on the model’s parameters but not on Ricco’s area and the shape of the fitted response profile). There was a small significant increase in the vertical Offset with eccentricity, which was reduced by accounting for convergence (**Figure 6 B and Table 2**).

**Figure 6.**
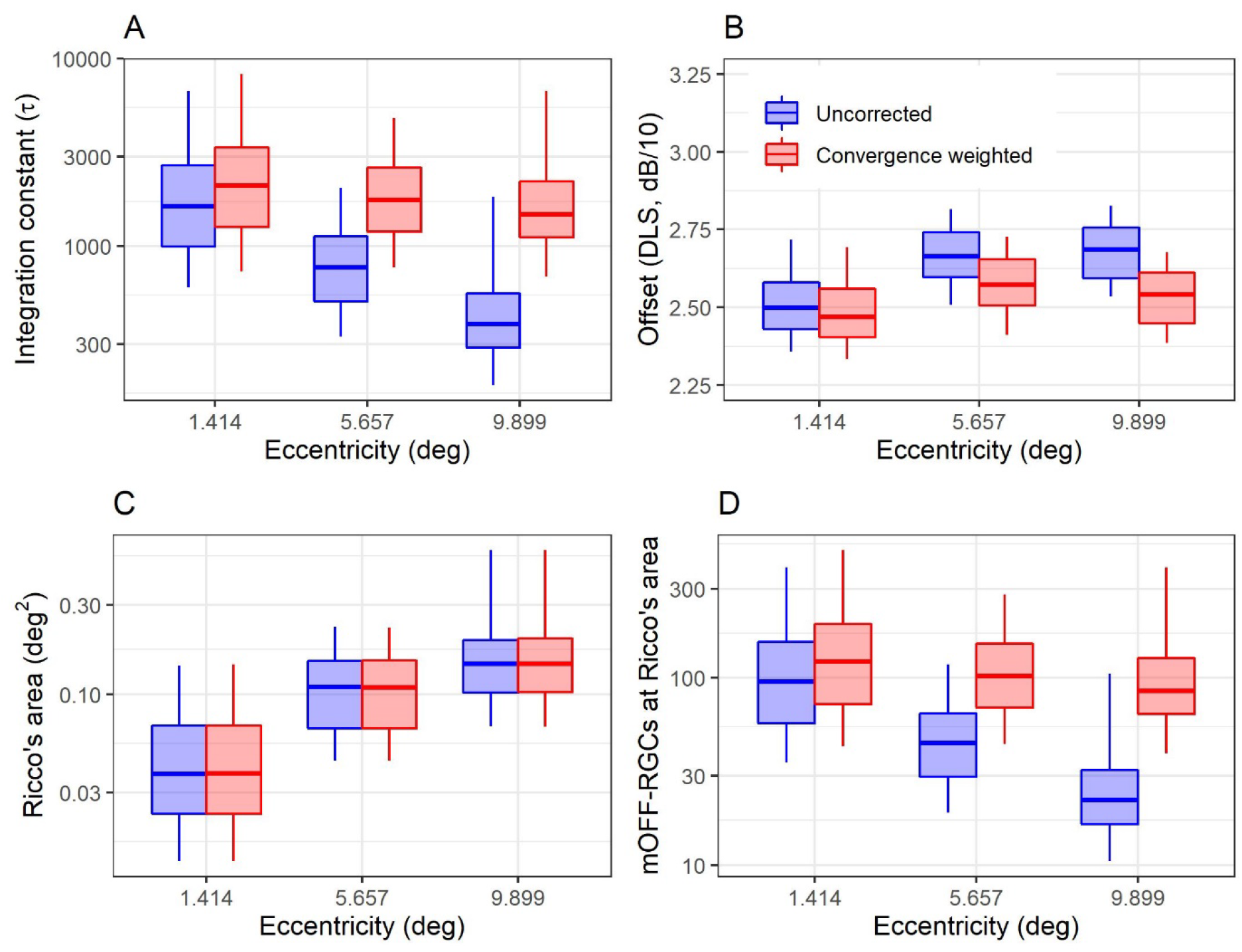
Boxplots of the different parameters and estimates derived from the model for spatial summation data. Note that the convergence weighted values in **(D)** are obtained by simply multiplying the uncorrected number of mOFF-RGCs at Ricco’s area by the convergence rate. The box encloses the interquartile range, the horizontal midline indicates the median and the error bars extend from the 5% to the 95% quantiles. The vertical axis is log_10_-spaced. RGC = Retinal Ganglion Cell

**Figure 7.**
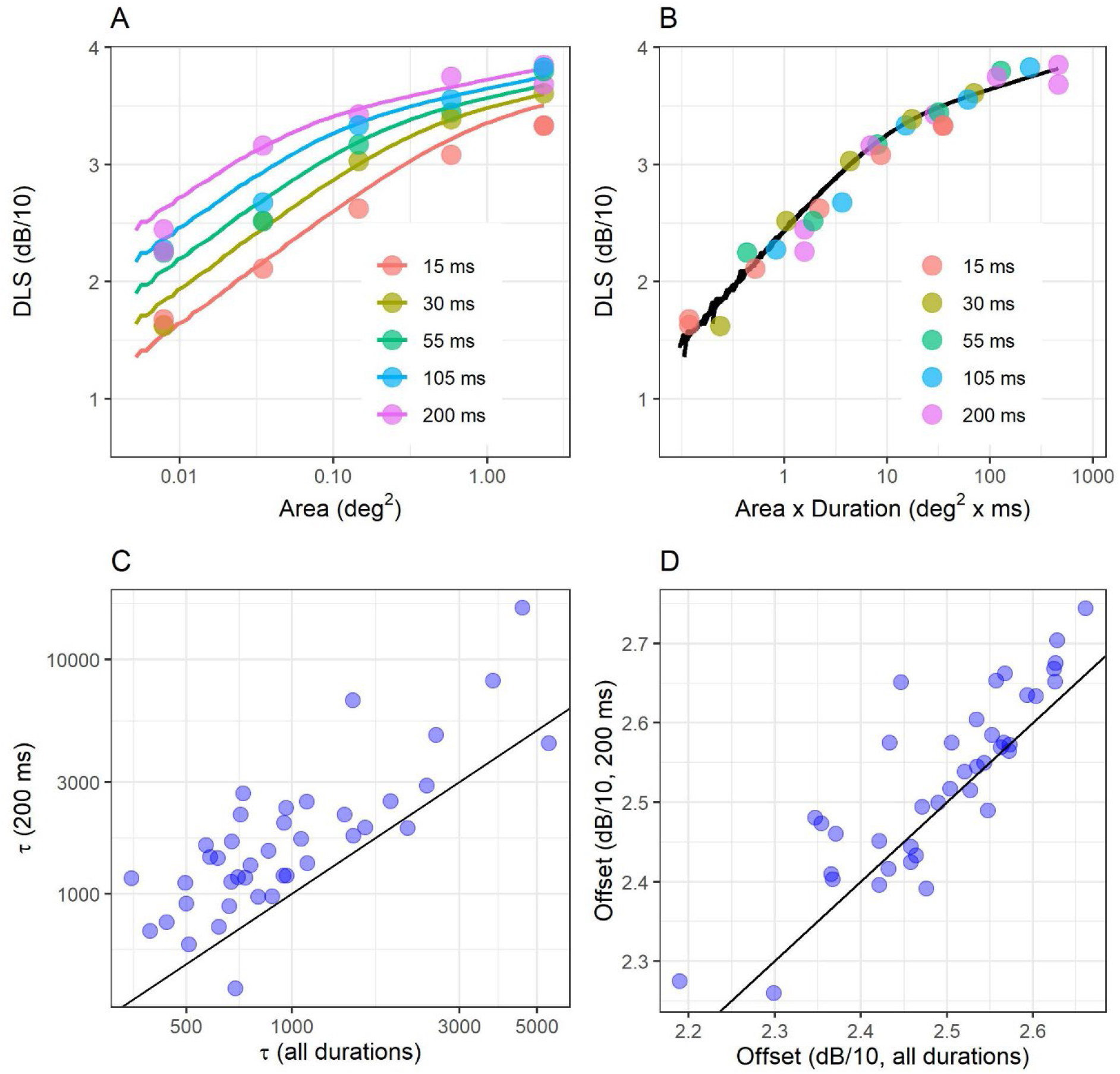
The two top panels show an example fit from one location in one subject, with the horizontal axis reporting the stimulus area **(A)** and the product of area and duration **(B)**. Correlation between the parameter estimates obtained by combining all durations and by only using data obtained with the 200 ms stimulus for the integration constant **(C)** and the offset **(D)**. The diagonal line indicates equivalence.

**Table 2.** Median [Interquartile Range] of the different outputs from the model fits. Comparisons were performed on log-transformed values but reported in linear scale (except for the Offset, which was tested and reported in log-scale and represents the shift in the relationship along the vertical axis). mOFF-RGC = midget OFF retinal ganglion cells. *Obtained by taking the product of Ricco’s area and local mOFF-RGC density; ^**†**^ Obtained by taking the product of Ricco’s area and local mOFF-RGC density scaled by retinal convergence.

|  |  | Eccentricity (degrees) |  |  | Comparisons |  |  |
| --- | --- | --- | --- | --- | --- | --- | --- |
|  |  | 1.414 (A) | 5.657 (B) | 9.899 (C) | A vs B | A vs C | B vs C |
| Uncorrected | $\tau$ ($\times 10^2$) | 16.42<br>[9.95, 27.06] | 7.7<br>[5.08, 11.3] | 3.84<br>[2.88, 5.6] | < 0.0001 | < 0.0001 | 0.0002 |
|  | Offset (dB/10) | 2.49<br>[2.43, 2.58] | 2.66<br>[2.6, 2.74] | 2.69<br>[2.59, 2.76] | < 0.0001 | < 0.0001 | 0.4975 |
|  | Ri'co's area (deg<sup>2</sup>) | 0.038<br>[0.023, 0.068] | 0.109<br>[0.066, 0.151] | 0.145<br>[0.102, 0.195] | < 0.0001 | < 0.0001 | 0.0158 |
|  | # mOFF-RGCs* | 95.2<br>[57.12, 156.06] | 44.82<br>[29.68, 64.64] | 22.26<br>[16.49, 32.22] | < 0.0001 | < 0.0001 | 0.0001 |
| Convergence weighted | $\tau$ ($\times 10^2$) | 21.08<br>[12.7, 33.64] | 17.65<br>[11.95, 26.24] | 14.76<br>[11.18, 22.14] | 0.9447 | 0.4457 | 0.9247 |
|  | Offset (dB/10) | 2.46<br>[2.4, 2.56] | 2.57<br>[2.51, 2.65] | 2.54<br>[2.45, 2.61] | < 0.0001 | 0.0301 | 0.0301 |
|  | Ri'co's area (deg<sup>2</sup>) | 0.038<br>[0.023, 0.068] | 0.109<br>[0.066, 0.152] | 0.145<br>[0.102, 0.198] | < 0.0001 | < 0.0001 | 0.0163 |
|  | # mOFF-RGCs<sup>†</sup> | 122.17<br>[72.59, 194.2] | 102.59<br>[69.25, 152.79] | 85.5<br>[64.27, 127.58] | 0.8699 | 0.4270 | 0.8699 |

##### Spatio-temporal summation

The same spatio-temporal model was used to analyse data from locations {±7; ±7} with all different combinations of stimulus sizes and durations. The data were collated to obtain a single estimate of the integration constant and accounting for retinal convergence. The global average RMSE for this fit was 1.67 ± 0.52 dB (Mean ± SD) and 1.40 ± 0.41 dB for the 200 ms stimuli. This can be compared to the 0.96 ± 0.35 dB average RMSE obtained from fitting the 200 ms data alone at the same eccentricity. For context, the root mean squared difference in sensitivity between the two repetitions of the retested combinations was 2.44 dB and the root mean squared deviation from the average of the two repetitions was 1.22 dB. An example of the calculation for one location in one subject is also shown (**Figure 7 A and B**). There was a strong correlation between the parameter estimates obtained by fitting data from all stimulus durations and 200 ms alone (previous section), at the same eccentricity (correlation coefficient: 0.79 for log_10_(τ) and 0.84 for the sensitivity offset, **Figure 7 C and D**). However, the two estimates appeared to have a consistent significant difference (p < 0.0001), approximately constant in log_10_-scale. The Median [Interquartile Range] was 34.65 [25.31, 56.04] x 10^2^ for the τ constant and 2.36 [2.31, 2.42] dB/10 for the offset. These values were both significantly smaller than those reported in **Table 2** for the same eccentricity (p < 0.0001 and p = 0.00298 respectively). Significant differences were also present for all the other parameters, including Ricco’s area and the number of mOFF-RGCs at Ricco’s area (all p < 0.0001). Numeric values of Ricco’s area and corresponding mOFF-RGC counts are reported in **Table 3** for all durations. Differences in Ricco’s areas between different durations were not tested as such differences are assumed by the model.

**Table 3.**
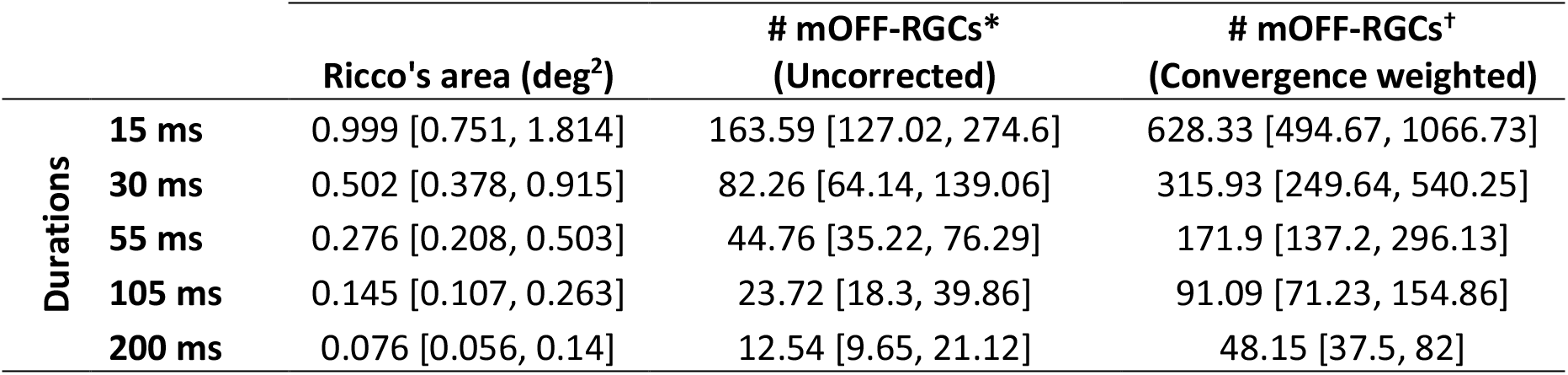
Median [Interquartile Range] of the different outputs from the model fits with the different stimulus durations. mOFF-RGC = midget OFF retinal ganglion cells. *Obtained by taking the product of Ricco’s area and local mOFF-RGC density; ^**†**^ Obtained by taking the product of Ricco’s area and local mOFF-RGC density scaled by retinal convergence.

#### Frequency of seen curves

We estimated the FoS curves for the four most extreme combinations of stimulus size and duration at locations {±7; ±7} using the MOCS data for five subjects. The results of the Bayesian fitting are shown in **Figure 9**. The FoS was modelled using the CDF of a Gaussian distribution. The average of the estimates for μ, σ, λ and γ are reported as **Supplementary material**.

**Figure 8.**
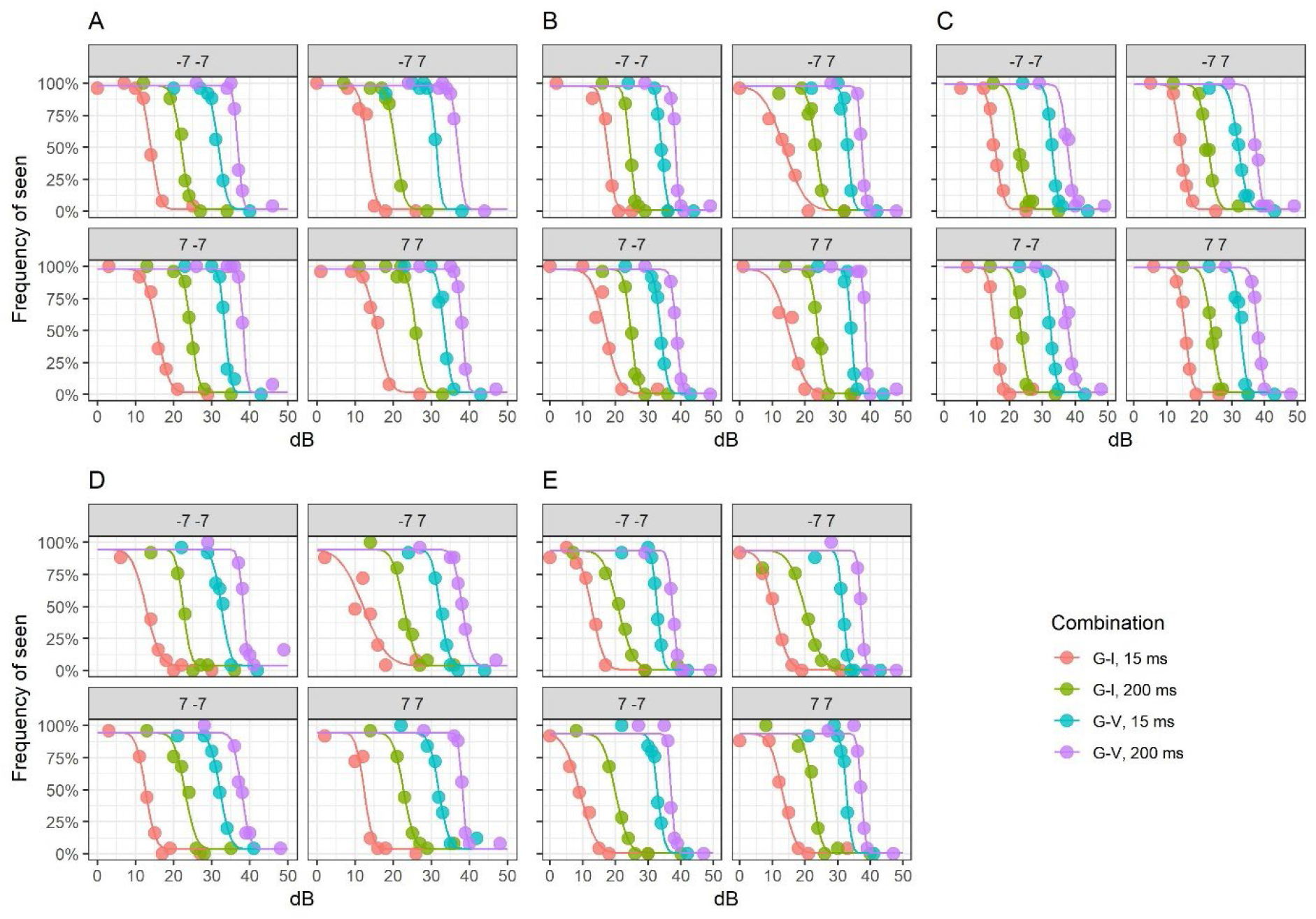
Fitted psychometric functions obtained from the Method of Constant Stimuli (MOCS) experiment at the four tested locations in five subjects with four different combinations of stimulus size and duration. Parameters are provided as **Supplementary material**.

**Figure 9.**
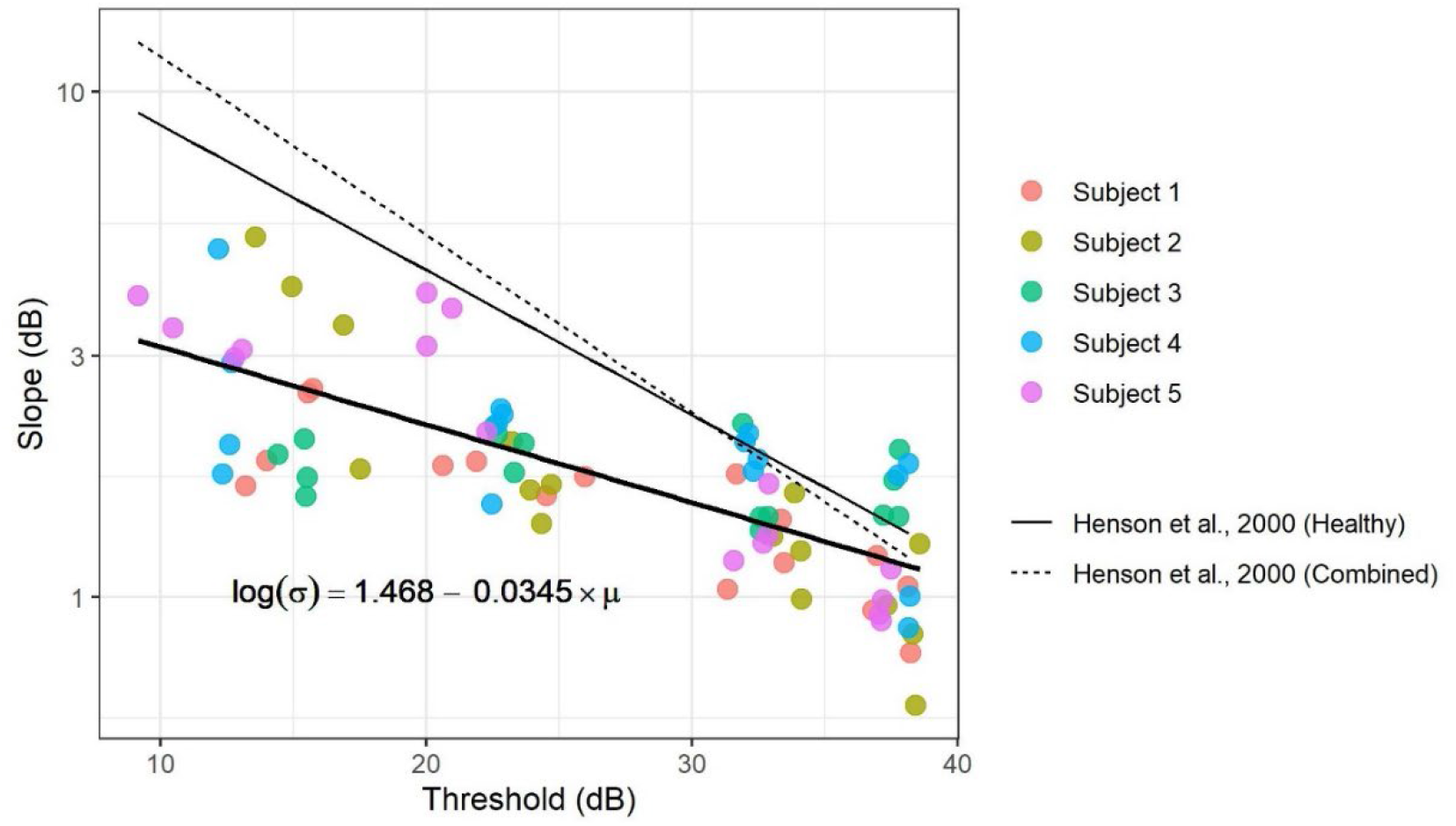
Relationship between the slope (σ) of the psychometric function and the 50% threshold (μ). The regression line is also reported. The relationship was statistically significant (p < 0.0001).

In general, there was a tendency for slopes (σ) to be shallower for conditions where sensitivity was lower (μ). This agrees with previous literature^45, 46^. **Figure 9** shows this relationship. Estimates from Henson et al.^45^ are also reported for comparison.

## Discussion

Constant integration of visual input has been regarded as a fundamental principle governing the perception of visual stimuli^18, 19^. However, the interaction of stimulus duration and size has been rarely and incompletely explored in perimetry^11^. Our data support constant input integration as a fundamental principle in perimetric response in healthy observers. Such a principle has translational value as it provides a simple framework for the interpretation and prediction of perimetric responses in healthy subjects and allows speculations on the expected changes from disease.

The first important result is the change in Ricco’s area with different stimulus durations. The size of Ricco’s area has often been interpreted considering cortical magnification^10^, linking the number of RGCs within Ricco’s area to the number of RGCs contacting V1 cells in the visual cortex. Such a line of reasoning seems however questionable if Ricco’s area can vary with stimulus duration, because duration would have no effect on the spatial extent of RGC-V1 connections. Rather, temporal and spatial summation appear to operate in concert to maintain a consistent behaviour in response to the same amount of visual input, be it from changes in stimulus size or duration. Fredericksen et al.^21^ also proposed a similar integration model in the context of motion detection, suggesting that spatio-temporal summation likely arise from diffuse cortical integration rather than specific temporal or spatial processes. Our model captures such a spatio-temporal interaction by only requiring the fitting of one parameter (the integration constant τ), while providing good predictions of the experimental results. Other models, while not specifically investigating the interaction between stimulus size and duration, also showed that the spatial scale of the visual system could be modelled independently of the underlying RGC density and their RFs using cortical filters with different spatial scales^1, 7^. Our model also decouples spatial summation from the extent of the retinal spatial filters (in this case, the extent of the DoG filter used to model RGCs’ responses). This has important implications for modelling the effect of disease that will be discussed later. It should be noted that other authors have proposed that these effects could be explained by a dynamic change in the “functional” receptive field size as a function of stimulus duration and background luminance^13^.

More realistically, this could correspond to a selection of cortical filters of different sizes for different stimulus characteristics or to the response envelope of multiple filters combined by probability summation whose sensitivity can be selectively changed by different stimulation conditions^1^. Further research is needed to understand how this would apply in the case of disease, such as RGC loss (see later). Such a mechanism is further explored in a dedicated paragraph in the **Appendix**.

The model described by Equations (5) and (6) can be modified to incorporate different impulse response functions. In this study, it was a simple capacitor equation, as this was deemed sufficient to model our data by fitting only two parameters. This is likely to be simplistic for many other applications. For example, our model does not include any response delay. Our results can be largely replicated with the monophasic response filter used by Gorea and Tyler^15^ and first described by Watson^47^. Such an impulse response can also be tuned to produce different critical durations by changing an integration constant, while keeping all the other parameters fixed. Using this impulse response only produced minimal differences (one example is provided as **Supplementary Material**). A drop in sensitivity has been shown for very long stimulus durations^48–50^ and modelled with a biphasic impulse response integrated over a limited time window^15^. Our stimuli would not be long enough for this to be evident. Our temporal integral in Equation (6) extends to infinity, similarly to Watson^14^. Gorea and Tyler^15^ highlighted the implausibility in this assumption, because an observer that integrates over an infinite time window will never make a decision to respond. A practical choice for our experiments would be to use the maximum time interval allowed for a response (1500 ms) as an integration window. However, this is so much longer than the longest stimulus (200 ms) that it would be practically equivalent to infinity.

It should be mentioned that both temporal and spatial summation, and contrast sensitivity in general, can be largely affected by background adaptation. For the background illumination used in this study (10 cd/m^2^) threshold behaviour should be close to Weber’s law at least for a G-III stimulus^51, 52^. Retinal illuminance can be reduced by media opacity (such as cataract) but this is likely to be negligible in a young healthy cohort. Pupil size can also affect retinal illuminance, especially if below 3 mm^51^, but the average pupil size in our cohort was 5.9 ± 0.8 mm.

The model can be used to investigate the effect of eccentricity on spatial summation. Our results show that Ricco’s area significantly increased with eccentricity, as expected^2–4^. However, this did not correspond to a constant number of mOFF-RGCs being stimulated, this number being comparably larger at smaller eccentricity. This is mirrored by the identical trend for the integration constant τ, indicating that more mOFF-RGCs need to be stimulated to achieve the same change in sensitivity closer to the fovea. Our results agree with Kwon and Liu^10^, who also observed a notable departure from a constant number of mOFF-RGCs at Ricco’s area and a trend with eccentricity. However, they concluded that this was likely a result of inaccuracies in the estimates of RGC density. We propose a different explanation: the trend in the number of mOFF-RGCs, and in the integration constant, appeared to be completely eliminated by weighting the contribution of each mOFF-RGC by the Cones:mOFF-RGC convergence ratio. This observation suggests that, much like the effect of change in stimulus duration, convergence can change the “contribution” provided by each RGC in terms of retinal input. Our model is able to account for this, because the contribution of each RGC can be weighted by its convergence rate prior to summation in Equation (6). Our experiments would not allow us to uncover a specific mechanism for this phenomenon. However, a reasonable hypothesis is that increased convergence could change the contrast gain determining the spiking rate of the RGC for a given level of contrast. For our analysis we considered one possible class of RGCs, mOFF-RGCs. This is important for our assumption of hexagonal tiling, because different classes of RGCs form independent and overlapping mosaics^53, 54^. mOFF-RGCs where chosen for comparison with recent literature^10^, and because they are the most prevalent type of RGCs in humans^31, 54^. However, previous literature showed that briefly flashed stimuli, such as those used in perimetry, might preferentially stimulate parasol RGCs^55^. We therefore confirmed our results by repeating all our analyses on spatial summation with the mosaic of parasol-OFF-RGCs (see **Supplementary Material**). The effect of eccentricity was much less pronounced, but there was still a significant difference in the number of stimulated RGC at Ricco’s area and in the integration constant between the smallest and the largest eccentricity. Interestingly, when weighted by convergence, the results were effectively identical to those obtained with the mOFF-RGC mosaic, because the higher convergence ratio for the parasol-OFF (P-OFF) mosaic effectively produced the same scaled input. It should be noted that there is no clear anatomical evidence of increased Cones:P-OFF-RGC convergence with eccentricity. However, this seems a reasonable assumption because the Cones: RGC ratio calculated from histology data^37, 56^ increases with eccentricity in a similar fashion for both the midget and parasol cells. The similarity between our results and those reported by Kwon and Liu^10^ should be interpreted with caution, because it can be explained by the fact that both our estimates and theirs were derived from those provided by Drasdo et al.^30, 31^, which are in turn based on a small histology dataset by Curcio and Allen^28^. Despite our attempt to improve precision by customising Drasdo’s estimates using individualised structural OCT data^30^, the results are unlikely to be greatly altered. Therefore, Kwon and Liu’s^10^ results cannot be considered a fully independent confirmation of our findings. Finally, it should be noted that the compensation of the effect of eccentricity with the convergence ratio might be coincidental and could be explained by other factors, such as optical aberrations. The effect of natural ocular optics on spatial summation in the parafoveal retina is debated^57–59^. In our model, we included the effect of optical aberrations and glare using the average MTF for the human eye proposed by Watson^38^: the data were fitted accounting for optical factors, but the summation curves were generated without the effect of optics. This was an attempt at estimating the pure neural contribution to spatial summation. However, the effect on the results largely depends on other assumptions within the model, particularly the choice of whether the summation in Equation (7) is taken over the signed or absolute value or the RGC response. Our choice of summing the signed contribution was based on some desirable properties of the model, particularly the perfect linear scaling of the response with the change in RGC density and filter size. This produced a very small effect from ocular optics, because the total power of the stimulus was simply spread over a larger area. Taking the summation over the absolute value instead produced a much greater effect (results reported in **Supplementary material**) because negative contributions from “inhibited” RGCs were transformed into positive contributions, greatly amplifying the effect of optical blur. Our choice of modelling produced an average change in Ricco’s area due to optical factors of 0.065 log_10_ units, which is very similar to the change measured by Tuten et al.^57^ with adaptive optics (AO). Taking the summation over the absolute value instead produced an average change of 0.69 log_10_ units, which is closer to what was reported by Dalimier and Dainity^58^ for similar experiments. Ultimately, a definitive answer to these questions could only be obtained by performing these same experiments with coupled AO corrected stimuli and imaging, so that accurate estimates of individual RGCs can be obtained and the effect of optical aberrations eliminated^60^.

Another important result is the effect of different stimulus durations and sizes on the shape of the psychometric function. In general, and in agreement with previous reports^45, 46^, we have found that the change in the slope of the psychometric function was largely explained by a change in sensitivity and was reasonably described by a log-linear relationship (**Figure 9**). This effect is indicative of the presence of multiplicative noise in the response^40^. However, it is difficult to identify the exact origin of such noise (quantal fluctuations; eye movements; noise from the instrument). This has however important implications, because it confirms that the increase in variability of perimetric responses with sensitivity is not uniquely linked to disease but can be replicated in healthy observers. The MOCS experiments were designed to replicate the simple detection task involved in perimetry, where observers are asked to continuously monitor the presence of a signal in sequential intervals. This can be modelled as a task with a variable observer-defined “criterion” (i.e. rate of false alarm or response bias)^61^. In our FoS curves, this bias is accounted for by estimating the guess rate as a lower asymptote (the γ term in Equation (2)). This framework is rooted in high-threshold theory and widely adopted in the field of perimetry^62^. It should be kept in mind that, under the alternative signal detection theory, the bias correction would be performed after z-score transformation and would require numerous catch trials to determine the individual response bias^61^. In our data, the response bias and lapse rate were estimated from the response to stimuli that were likely to be much above or below the 50% threshold (as determined using a pilot using QUEST+ to estimate threshold and psychometric function slope) and all participants were encouraged to maintain a low false-alarm rate during the experiments. Both the guess and lapse rates were very close to 0 and are therefore unlikely to have greatly affected the estimates of the psychometric function.

Our choice of placing our testing locations along the diagonals limits our ability to appreciate the previously reported dissociation in between ganglion cell number and perimetric sensitivity in nasal visual field^63^. We however found a significantly smaller number of mOFF-RGCs within Ricco’s area for the nasal locations, indicating a smaller spatial scale compared to temporal locations (p = 0.005). This comparison was performed for the log_10_-RGC number with a linear mixed model using the hemifield as a fixed effect and the eccentricity as a random effect, nested within the subject, to perform a paired same-eccentricity comparison.

It is interesting to consider the implications of our results and modelling approach for the interpretation of changes observed in disease. Redmond et al.^9^ have demonstrated an increase in Ricco’s area in patients with glaucoma, which could be accounted for by a shift of the summation curves along the horizontal axis (stimulus size). According to some models^7, 10^, such a change could only occur by scaling the spatial filters to increase spatial convergence (equivalent to changing the cortical magnification factor), which would imply some sort of “restructuring” of either the pooling mechanism (for example the spatial extent of RGC-V1 connections) or an enlargement of RGCs’ RFs. The latter seems implausible, because most histologic studies have shown dendritic pruning and shrinkage^64^, which would imply smaller RGCs’ RFs. The first hypothesis also lacks solid support from experiments: Wang et al.^65^ observed changes in the cortical magnification factor in patients with glaucoma tested with functional magnetic resonance imaging; such changes, however, are indicative of increased retina-V1 divergence, and therefore do not clearly support the hypothesis of an increased magnification factor. Our model makes no such assumptions. Instead, the change in Ricco’s area is a consequence of the reduction in retinal input owing to a loss of RGCs in glaucoma. In **Figure 10 A**, data from healthy participants in Redmond et al.^9^ were fitted with our model, assuming a mosaic of mOFF-RGCs with density estimated from Drasdo et al.^30, 31^. The mosaic was then randomly degraded to achieve 73% RGC loss, equivalent to the reported proportional average change in Ricco’s area. The figure plots the average response of 100 randomly degraded mosaics. The model correctly predicted a horizontal shift of the curve, in agreement with the data. A horizontal shift in the response could also be explained by RGC loss preferentially affecting higher frequency cortical filters, whose loss in sensitivity might determine a horizontal shift of their probability summation envelope^1^. Our model also predicts that temporal summation curves can be equated between healthy and glaucoma by appropriately scaling stimulus size. This is shown in **Figure 10 B**, for the same mosaics simulated in **Figure 10 A**. Mulholland et al.^11^ provided experimental evidence that using Ricco-scaled stimuli could reduce the difference in temporal summation observed between glaucoma patients and healthy controls with G-III stimuli, although some residual differences were still present. This is further proof of the interaction between stimulus size and duration. However, more research is needed to fully characterise such an interaction in glaucoma. Finally, our model also predicts changes in spatial and temporal summation with photoreceptor loss, such as from diseases of the external retina. However, studies investigating this with perimetric stimuli are still lacking and will need further research.

**Figure 10.**
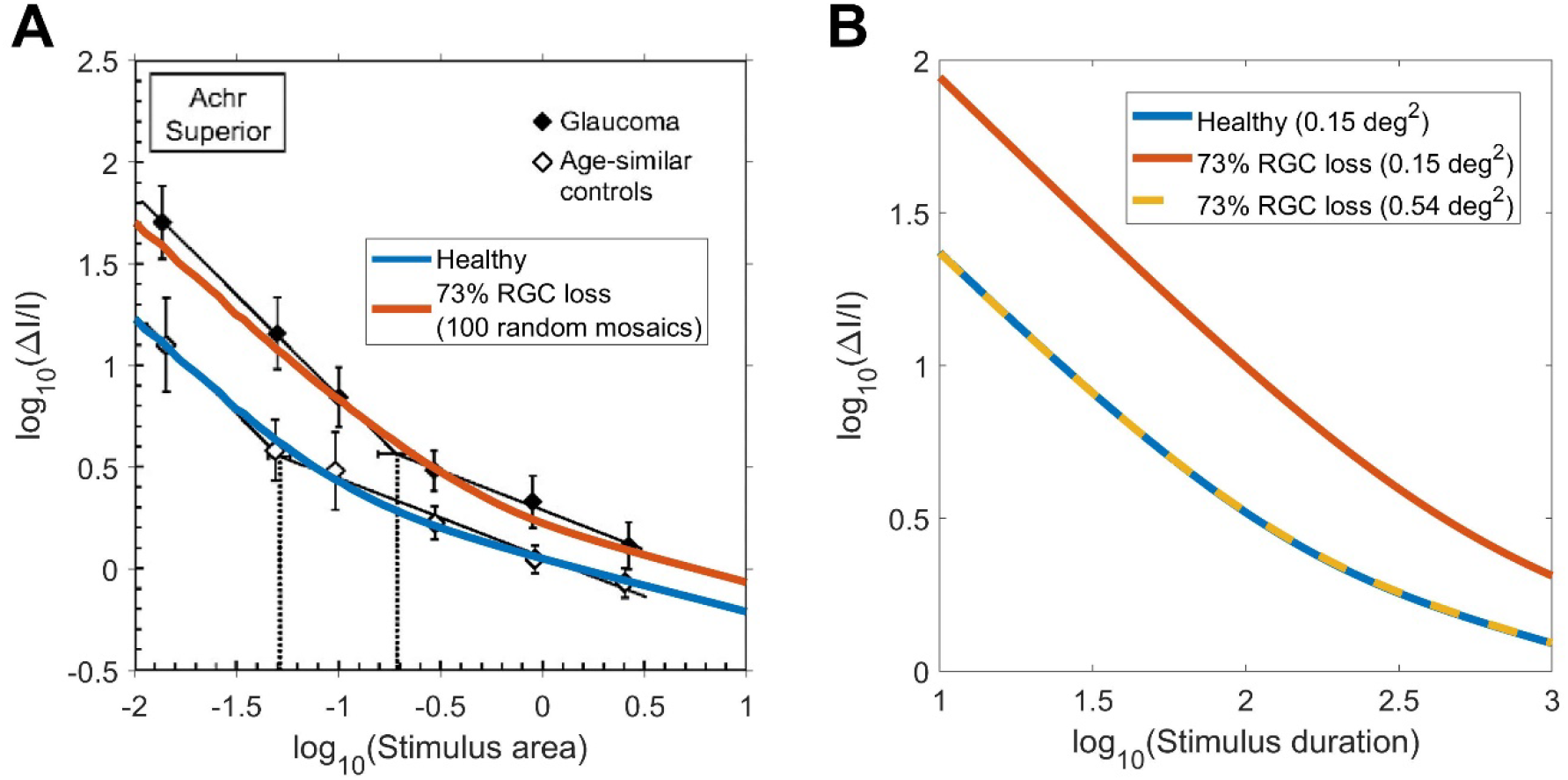
A) Change in Ricco’s area in patients with glaucoma compared to age-matched controls, adapted from Redmond et al.^9^. B) Temporal summation curves can be equated when RGC loss is compensated by an increase in the stimulus size. RGC = retinal ganglion cells.

Other questions remain, particularly pertaining to the systematic difference between the estimates of the model parameters obtained with 200 ms stimuli only or with all stimulus durations combined. Small inaccuracies in the delivery of the stimulus might produce variations in the intended durations, skewing the results of the combined analysis. Another consideration is that our model, despite describing most of the variability in the data, might not be capturing all aspects of the effect of stimulus duration on sensitivity. In fact, the model was not meant to be a complete description of the psychophysical response to all the features of the stimulus, but rather aimed at providing a coherent framework to explain important experimental observations from the data that are often neglected by other modelling attempts.

## Conclusions

We show that the amount of total retinal input can account for most of the characteristic of spatio-temporal summation with perimetric stimuli in healthy observers, including the effect of eccentricity. This could have important implications for the interpretation and design of perimetric examinations in diseased eyes as well as providing a framework for analysing spatio-temporal integration in heathy observers.

## Appendix

### MOCS fitting

MOCS data were fitted using a Bayesian hierarchical model. Each subject was fitted independently. The four combinations of stimulus size and duration were modelled as fixed effect factors on the parameters μ and σ of the psychometric function, as defined in Equation (2). The parameters for each stimulus combination are denoted as μ_c_ and σ_c_ The four locations were modelled as hierarchical random effects on μ_c_ and σ_c_, with no correlations between the two parameters. The lapse rate (λ) and guess rate (γ) were modelled as global parameters for the whole test. The response (yes/no) was modelled as binomial process with 25 trials. Following Prins et al.^27^, the prior distribution for the mean of the parameter μ_c_ and σ_c_ for each stimulus combination was a non-informative normal distribution with a standard deviation of 30 dB. The prior distribution for the variance of the parameters μ_c_ and σ_c_ for each stimulus combination was a non-informative uniform distribution between 0 and 1000 dB. The random effects for each location were modelled as a normal distribution with mean μ_c_ and standard deviation σ_c_. The prior distribution was linked to the parameter σ_c_ via a logarithmic function. The parameters γ and λ were non-hierarchical and had a Beta prior distribution with shape parameters 2 and 50.

The model was fitted by running two parallel MCMCs in Just Another Gibbs Sampler (JAGS)^66^. We used 5,000 burn-in iterations. After that, the model was run for 10,000 iterations. All parameters achieved a Gelman-Rubin diagnostic < 1.2^67^.

### Mosaic arrangement for computation

**Appendix Figure 1.**
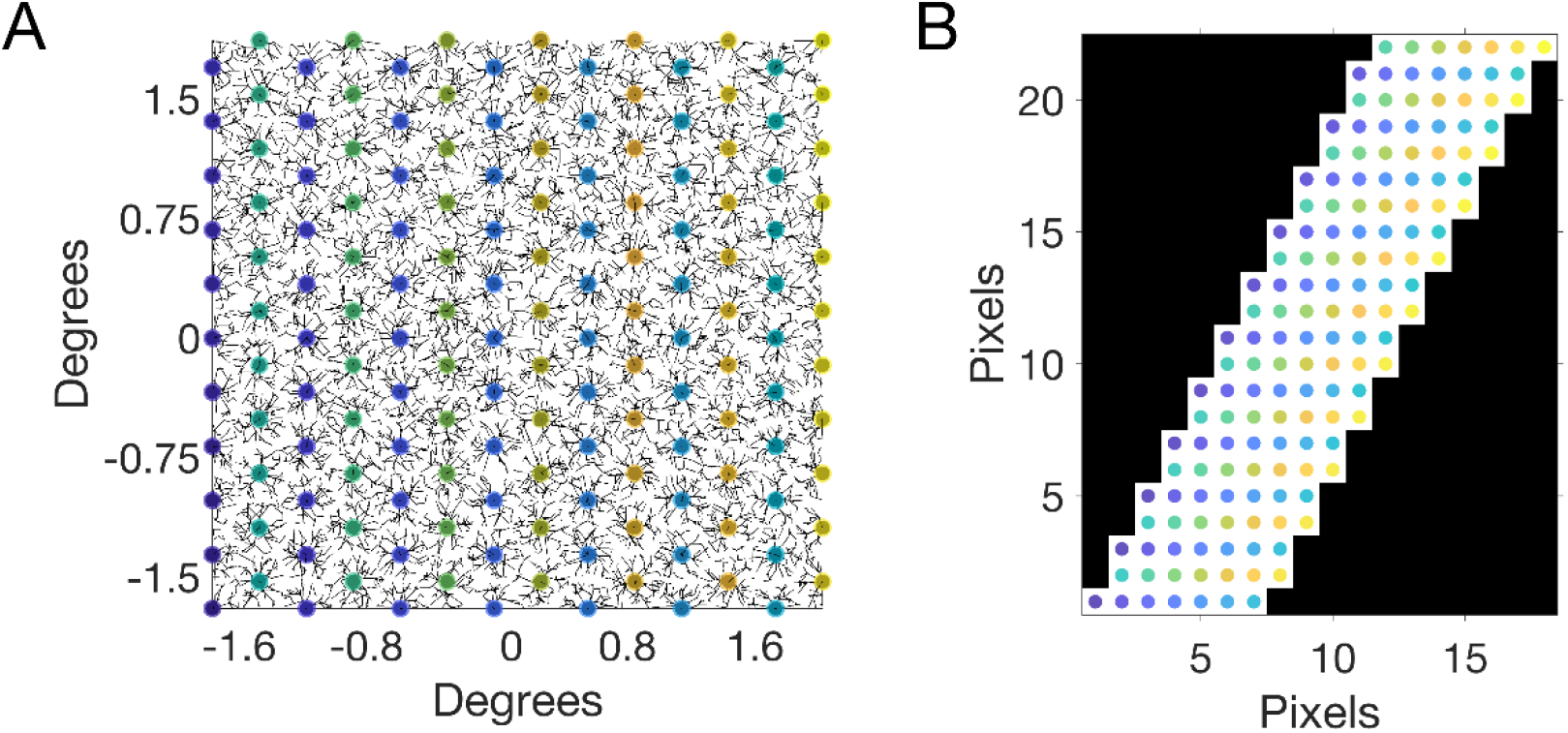
Example of how a cellular mosaic (RGCs in this case, left panel) is rearranged into a regular matrix with anisotropic spacing (right panel).

### A multiscale filter hypothesis for spatio-temporal integration

Many possible mechanisms could replicate the interaction between spatial and temporal summation reported in the literature and observed in our experiments. Our modelling approach is able to capture this aspect of the response. Nevertheless, it is useful to hypothesize how such an interaction could be implemented in the visual system. Glezer et al.^13^ proposed that this be achieved by a dynamic change in the “functional” RGC-RFs in the retina in response to changes in stimulation conditions, such as background illumination. However, there is no clear evidence of such a change occurring in the retina. Furthermore, Glezer et al.^13^ proposed such changes to occur through alterations in the weighting of the centre and surround of centre-surround receptive fields. Despite this, Ricco’s area was observed to alter in response to glaucomatous RGC loss in glaucoma patients^9^ and background luminance in healthy subjects^68^ in the s-cone pathway, in which a centre-surround receptive field organisation is absent^69^. A more reasonable hypothesis, that fits more closely with experimental observations, is that a set of spatial cortical filters exist and can be optimally selected based on the amount of retinal input. **Appendix Figure 2** shows a hypothetical response of an array of cortical neurons employing a biphasic first gaussian derivative filter (D1) with a gaussian envelope. This filter was chosen because it produces a smooth monotonic spatial summation curve, as shown by Pan and Swanson^1^. Note that the locations of the cortical neurons in the schematic indicate their projection into the visual space, rather than their anatomical arrangement in the visual cortex. In the schematic, selecting a larger filter corresponds to selecting a sparser mosaic of cortical neurons, since the extent of the filter is scaled with the inter-cellular spacing. This is equivalent to proportionally scaling the same mosaic. As expected, the summation curves with larger filters are shifted along the horizontal axis towards larger stimulus sizes. These mosaics can be obtained by selecting subsets of neurons from the same array (as in this example) or be constituted of separate sets of neurons. It should be mentioned that the summation curves produced by a more realistic implementation of this model (with cortical neurons sampling the response of RGCs with static RF sizes) would largely reproduce this behaviour but would not be an exact horizontal translation of the same response (see later).

**Appendix Figure 2.**
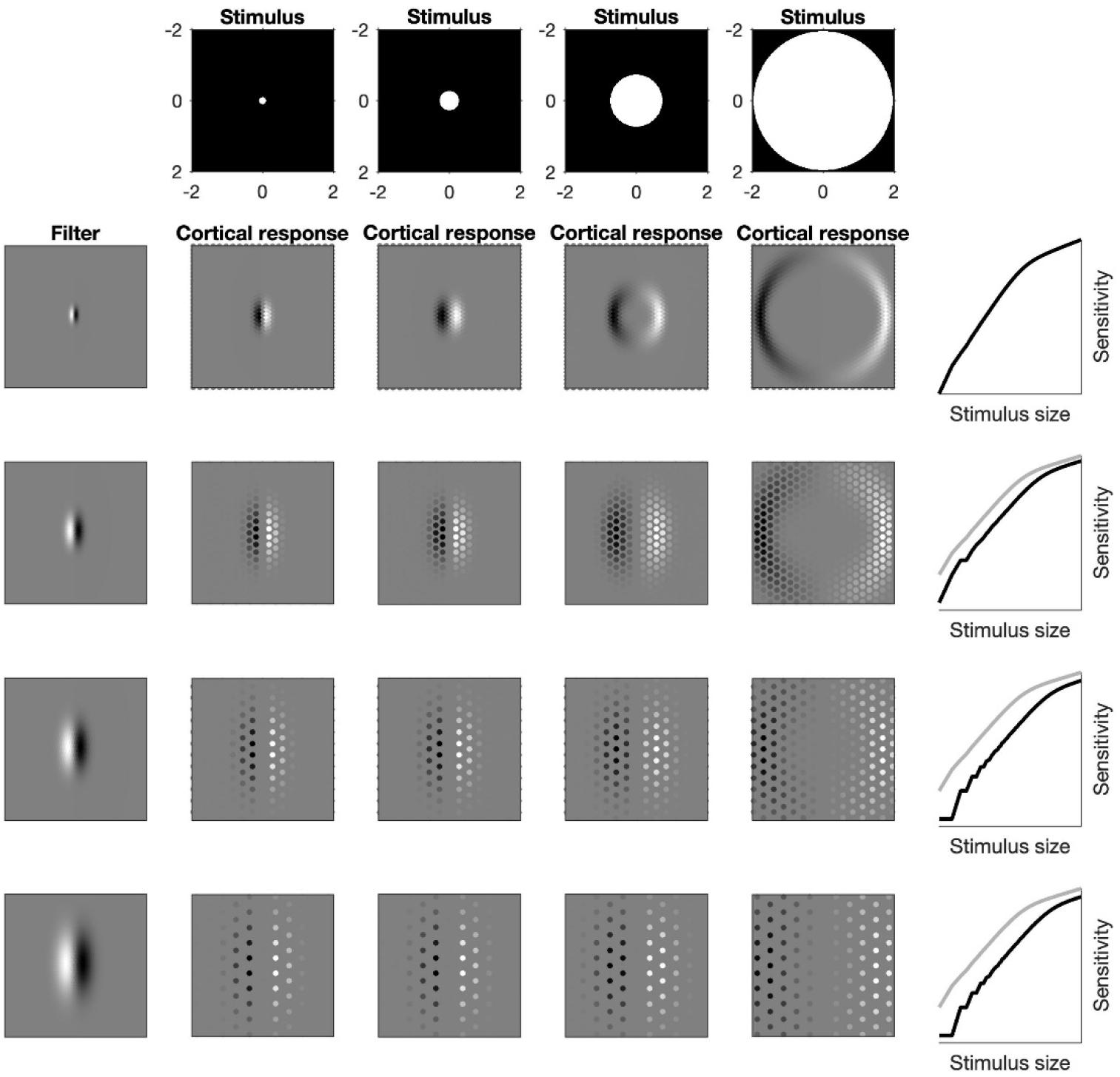
Example of how a change in scale results in a horizontal translation of the spatial summation curve. The cortical response is obtained by convolution of the spatial filter (left column) with the stimulus (top row). The summation curves (right column) are calculated as in Pan and Swanson^1^ with an exponent of 2. For this specific spatial filter, this corresponds to a partial summation slope of 0.25 in the log_10_ – log_10_ plot, the same as in our model. The spatial summation curve with the smallest filter is shown in grey for reference.

The change in spatial scale with different stimulus durations can therefore be replicated by a horizontal shift (in log_10_ – log_10_ coordinates) of the same template response by an amount equivalent to the log_10_ change in duration. Note that the selection of the filter scale does not need to depend solely on the stimulus duration, but more generically on the retinal input, to include the effect of Cones:RGC convergence, duration, background illumination or, for example, RGC loss in disease. For the sake of simplicity, everything except duration was held constant for these calculations. The combined effect is best represented by a summation surface, shown in **Appendix Figure 3**. In the figure, three summation curves are isolated by cutting through the surface at different stimulus durations and correspond to using a different filter scale. Importantly, temporal summation responses can be obtained by cutting through the surface along the orthogonal (duration) axis. Because the surface is obtained by proportionally translating the same spatial summation curve, temporal summation responses also follow the same template curve, proportionally shifted with different stimulus sizes. This would produce the same results obtained with our more generic input summation model. With this interpretation, although a strict retina-V1 convergence cannot be defined, testing in partial summation condition (i.e. long stimulus durations and high background illumination) would allow the calculation of the smallest possible spatial scale for a given retinal location.

**Appendix Figure 3.**
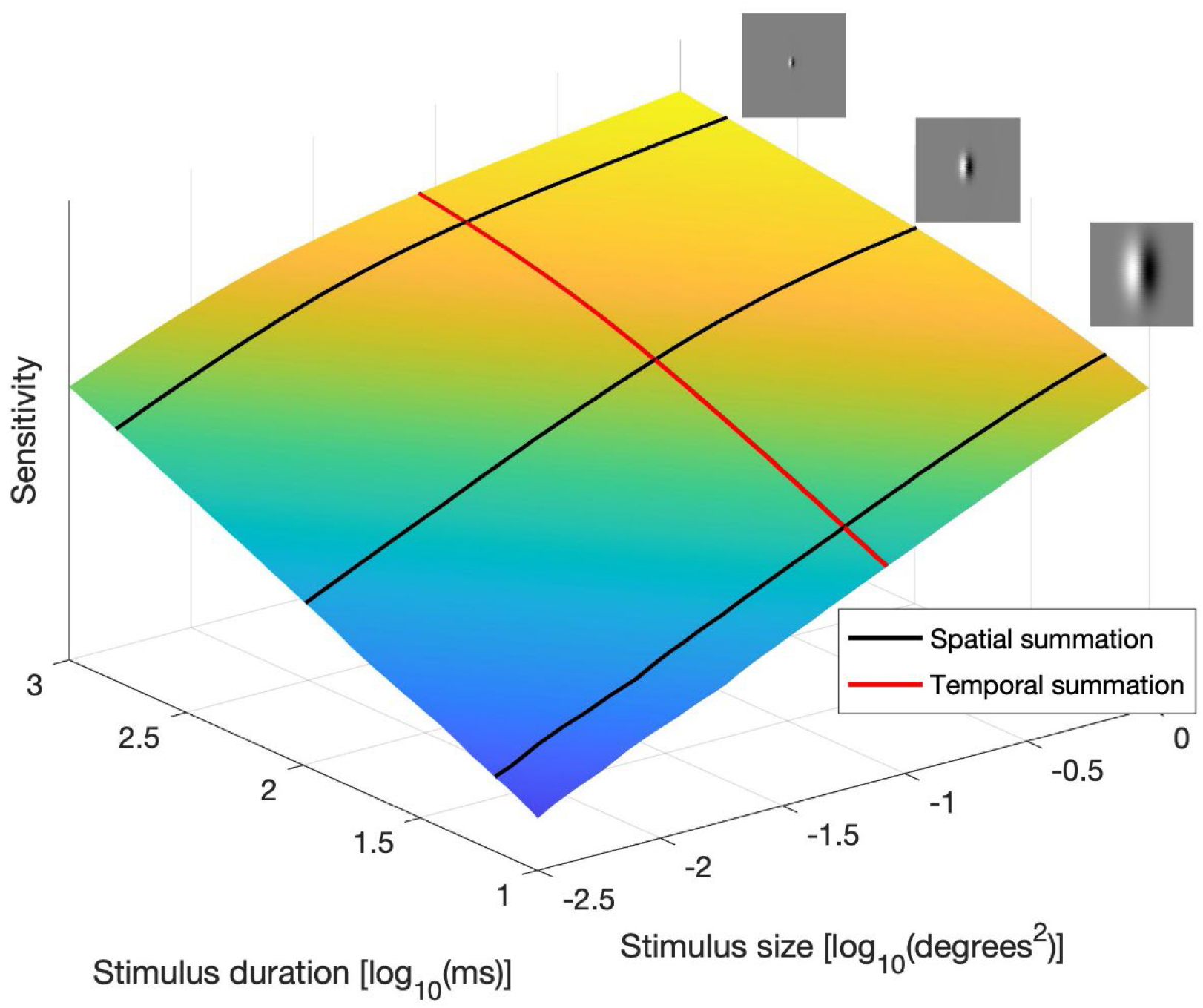
Spatio-temporal response surface, obtained by shifting the spatial summation curve by an amount equivalent to changes in duration, in log_10_ scale. Spatial and temporal summation curves are shifted versions of the same curve and can be obtained by cutting through the surface along different axes. The small insets show the change in the spatial filter for three different stimulus durations, producing the spatial summation curves identified by the black profiles.

Another possibility, proposed by Pan and Swanson^1^, is that different stimulus features, such as adaptation state and stimulus duration, might alter the relative sensitivity of individual filters and change the combined response “envelope” obtained through probability summation. For simplicity, we demonstrate this concept in **Appendix Figure 4** by selectively combining the response of filters with progressively smaller spatial scales. The resulting response envelope is a simple translation of the same curve.

**Appendix Figure 4.**
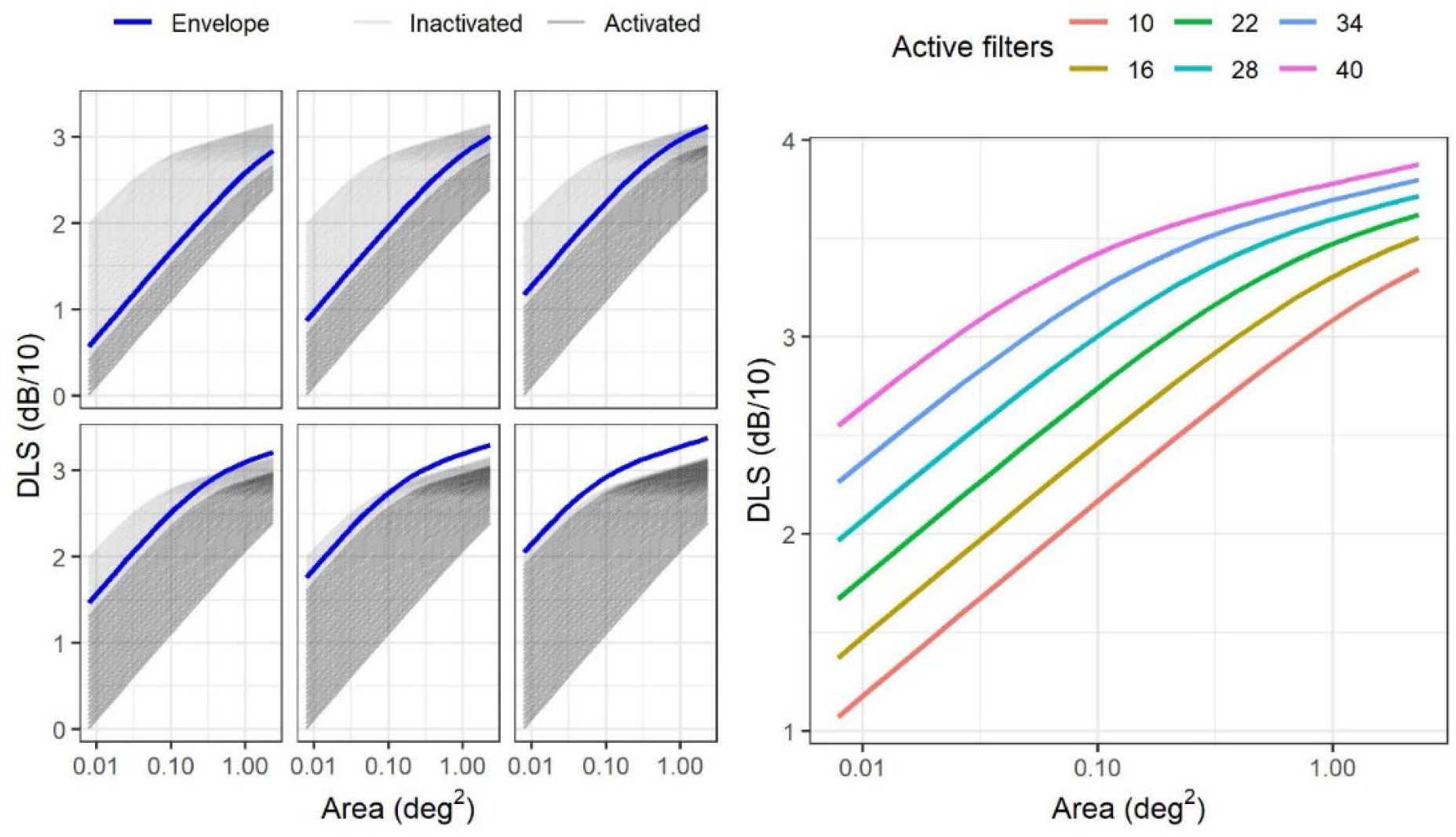
The blue lines in the left panels represent the envelope of the combine responses of the cortical filters whose responses are shown as black likes (inactive filters are in light grey). The right panel reports the same response envelopes, color-coded according to the number of hypothetical filters combined to generate the response.

We finally implemented a more realistic two-stage version^70^ of the cortical pooling model presented in **Appendix Figure 2**, where an array of cortical cells would sample the response of an array of RGCs like the one used in our main model. The cortical cell array was the same as the RGC array but used a D1 filter as their receptive field. **Appendix Figure 5** shows the responses produced by both the multiscale filters and the combination envelope. These largely replicate our experimental results (horizontal translation of the same response), with some small changes at different scales introduced by the fact that the size of the RGCs did not scale with the chosen cortical filter. Like in Swanson et al.^70^, this modelling exercise shows that Ricco’s area can be entirely determined by cortical filters without changing the RGC density or the size of the RGC-RF.

**Appendix Figure 5.**
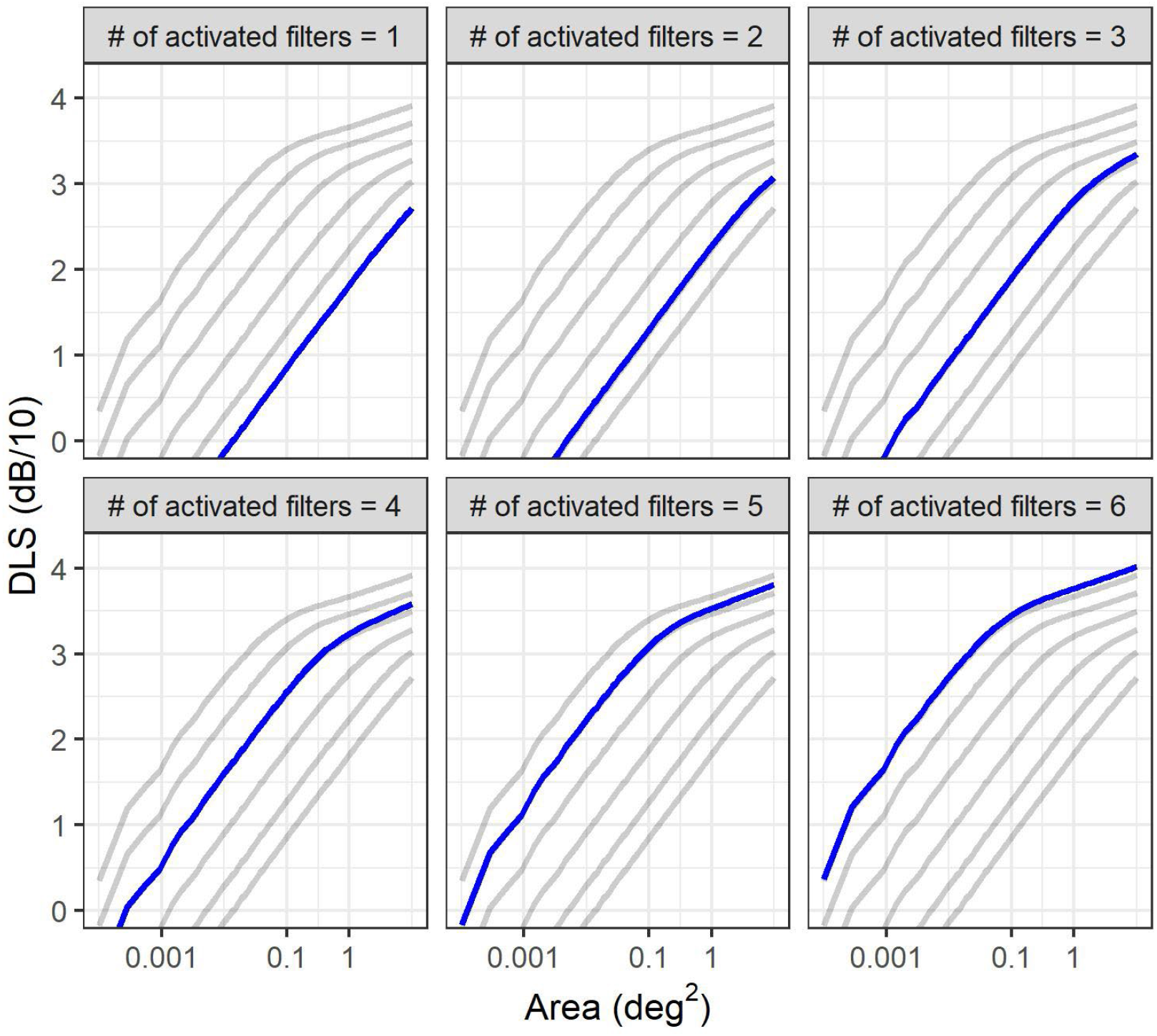
Replication of the model in **Appendix Figure 2** using a two-stage model. The individual filter responses at different scales are reported in grey. The blue line represents the response envelope obtained by combining, through probability summation, the responses of filters with progressively smaller spatial scales. For example, in the top-right panel, the envelope is obtained by combining the responses of the filters with the three largest scales, while excluding the remaining 3 with a smaller spatial scale.

