## Supplemental appendix for "Spatio-temporal summation of perimetric stimuli in healthy observers"

### Spatial summation with parasol OFF retinal ganglion cells

| | | Eccentricity (degrees) | | | Comparisons | | |
|---|---|---|---|---|---|---|---|
| | | 1.414 (A) | 5.657 (B) | 9.899 (C) | A vs B | A vs C | B vs C |
| Uncorrected | $\tau$ (x $10^2$) | 3.07 [1.81, 4.89] | 2.39 [1.6, 3.5] | 1.63 [1.22, 2.53] | 0.2164 | 0.0074 | 0.1325 |
| | Offset (dB/10) | 2.67 [2.61, 2.76] | 2.79 [2.72, 2.88] | 2.78 [2.69, 2.84] | < 0.0001 | 0.0002 | 0.2798 |
| | Ricco's area (deg$^2$) | 0.039 [0.023, 0.067] | 0.111 [0.067, 0.149] | 0.146 [0.104, 0.191] | < 0.0001 | < 0.0001 | 0.0132 |
| | # P-OFF-RGCs* | 18.08 [10.5, 28.29] | 13.67 [9.23, 20.26] | 9.34 [6.85, 14.2] | 0.1843 | 0.0043 | 0.1104 |
| Convergence weighted | $\tau$ (x $10^2$) | 21.54 [12.95, 33.6] | 18.09 [12.31, 26.41] | 14.55 [11.21, 23.78] | 0.9999 | 0.5822 | 0. 9999 |
| | Offset (dB/10) | 2.46 [2.4, 2.56] | 2.57 [2.5, 2.66] | 2.54 [2.44, 2.61] | < 0.0001 | 0.0406 | 0.0406 |
| | Ricco's area (deg$^2$) | 0.039 [0.024, 0.067] | 0.11 [0.067, 0.152] | 0.145 [0.104, 0.198] | < 0.0001 | < 0.0001 | 0.0134 |
| | # P-OFF-RGCs† | 126.25 [74.27, 194.57] | 103.17 [69.98, 152.93] | 83.29 [62.86, 133.73] | 0.9241 | 0.4364 | 0.9241 |

**Table S1**. Median [Interquartile Range] of the different outputs from the model fits. Comparisons were performed on log-transformed values but reported in linear scale (except for the Offset, which was tested and reported in log-scale). P-OFF-RGC = parasol OFF retinal ganglion cells. *Obtained by taking the product of Ricco's area and local P-OFF-RGC density; † Obtained by taking the product of Ricco's area and local P-OFF-RGC density scaled by retinal convergence.

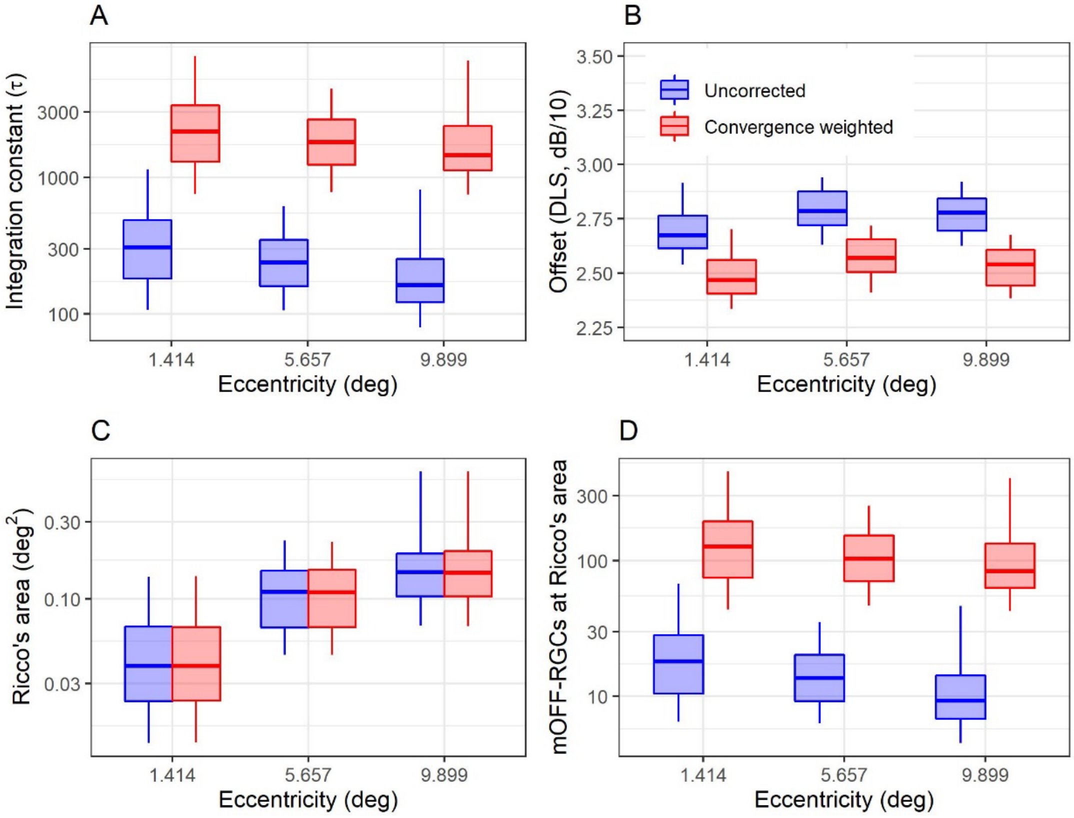


**Figure S1.** Box-plots of the different parameters and estimates derived from the model for spatial summation data. Note that the convergence weighted values in **(D)** are obtained by simply multiplying the uncorrected number of parasol OFF-RGCs at Ricco's area by the convergence rate. The box encloses the interquartile range, the horizontal midline indicates the median and the error bars extend from the 5% to the 95% quantiles. The vertical axis is $log_{10}$-spaced. RGC = Retinal Ganglion Cell

### Effect of optical factor compensation

| | | Eccentricity (degrees) | | | Comparisons | | |
|---|---|---|---|---|---|---|---|
| | | 1.414 (A) | 5.657 (B) | 9.899 (C) | A vs B | A vs C | B vs C |
| Uncorrected | τ (x $10^2$) | 24.72 [16.74, 42.77] | 16.95 [12.00, 26.78] | 9.14 [6.66, 13.68] | 0.0135 | < 0.0001 | 0.0016 |
| | Offset (dB/10) | 2.47 [2.41, 2.57] | 2.61 [2.55, 2.71] | 2.63 [2.54, 2.70] | < 0.0001 | < 0.0001 | 0.7222 |
| | Ricco's area (deg$^2$) | 0.027 [0.017, 0.050] | 0.112 [0.067, 0.160] | 0.156 [0.101, 0.211] | < 0.0001 | < 0.0001 | 0.0080 |
| | # mOFF-RGCs* | 63.54 [41.79, 115.41] | 42.9 [30.09, 69.55] | 23.58 [17.09, 35.93] | 0.0110 | < 0.0001 | 0.0040 |
| Convergence weighted | τ (x $10^2$) | 30.67 [20.56, 53.60] | 40.98 [28.01, 62.18] | 35.76 [25.96, 54.17] | 0.4547 | 0.5955 | 0.6915 |
| | Offset (dB/10) | 2.44 [2.39, 2.55] | 2.52 [2.46, 2.62] | 2.48 [2.40, 2.56] | 0.0134 | 0.6821 | 0.0283 |
| | Ricco's area (deg$^2$) | 0.027 [0.017, 0.050] | 0.112 [0.07, 0.161] | 0.155 [0.102, 0.212] | < 0.0001 | < 0.0001 | 0.0089 |
| | # mOFF-RGCs$^{\dagger}$ | 78.79 [51.62, 143.94] | 104.15 [69.84, 161.06] | 92.28 [65.06, 142.37] | 0.7705 | 0.8026 | 0.8026 |

**Table S2**. Median [Interquartile Range] of the different outputs from the model fits by taking the summation over the module (absolute value) of the RGC. Comparisons were performed on log-transformed values but reported in linear scale (except for the Offset, which was tested and reported in log-scale). mOFF-RGC = midget OFF retinal ganglion cells. *Obtained by taking the product of Ricco's area and local, mOFF-RGC density; $^{\dagger}$ Obtained by taking the product of Ricco's area and local mOFF-RGC density scaled by retinal convergence.

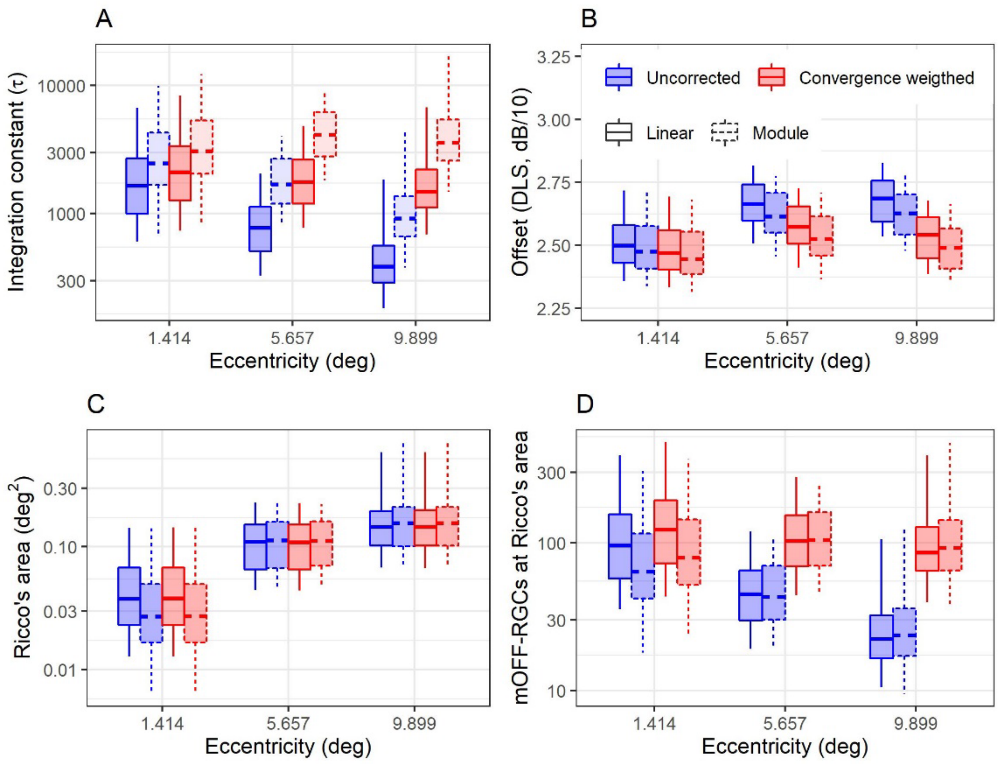


**Figure S2.** Box-plots of the different parameters and estimates derived from the model for spatial summation data. The shading indicates whether the summation in Equation (6) in the text was taken over the absolute value (module) or the signed (linear) RGC input. The box encloses the interquartile range, the horizontal midline indicates the median and the error bars extend from the 5% to the 95% quantiles. The vertical axis is $log_{10}$-spaced. RGC = Retinal Ganglion Cell; OF = Optical factors (average modulation transfer function of the eye)

### Average parameters for the psychometric functions

| Stimulus | Location {X, Y} | μ, σ (dB) | | | | |
|---|---|---|---|---|---|---|
| | | **Subject 1** | **Subject 2** | **Subject 3** | **Subject 4** | **Subject 5** |
| **G-I, 15 ms** | **{-7, -7}** | 13.99, 1.86 | 17.52, 1.79 | 15.43, 2.06 | 12.68, 2.90 | 12.79, 2.97 |
| | **{-7, 7}** | 13.22, 1.66 | 13.59, 5.15 | 14.41, 1.92 | 12.17, 4.89 | 10.46, 3.41 |
| | **{7, -7}** | 15.55, 2.54 | 16.88, 3.45 | 15.48, 1.58 | 12.60, 2.00 | 9.16, 3.94 |
| | **{7, 7}** | 15.72, 2.58 | 14.95, 4.12 | 15.52, 1.72 | 12.35, 1.75 | 13.08, 3.09 |
| **G-I, 200 ms** | **{-7, -7}** | 21.90, 1.86 | 24.35, 1.4 | 22.67, 2.10 | 22.46, 1.53 | 20.97, 3.73 |
| | **{-7, 7}** | 20.63, 1.82 | 23.24, 2.03 | 22.56, 2.18 | 22.80, 2.36 | 20.03, 3.99 |
| | **{7, -7}** | 24.54, 1.59 | 24.70, 1.67 | 23.32, 1.76 | 22.90, 2.29 | 20.03, 3.13 |
| | **{7, 7}** | 25.96, 1.73 | 23.92, 1.63 | 23.68, 2.02 | 22.69, 2.20 | 22.28, 2.12 |
| **G-V, 15 ms** | **{-7, -7}** | 31.69, 1.75 | 34.09, 1.24 | 32.87, 1.44 | 32.49, 1.88 | 32.86, 1.33 |
| | **{-7, 7}** | 31.34, 1.04 | 33.06, 1.32 | 31.91, 2.20 | 32.30, 1.77 | 31.56, 1.18 |
| | **{7, -7}** | 33.47, 1.17 | 33.86, 1.61 | 32.56, 1.35 | 32.12, 2.11 | 32.90, 1.68 |
| | **{7, 7}** | 33.37, 1.43 | 34.12, 0.99 | 32.57, 1.44 | 32.00, 2.03 | 32.67, 1.28 |
| **G-V, 200 ms** | **{-7, -7}** | 36.81, 0.94 | 38.31, 0.85 | 37.61, 1.70 | 38.21, 1.00 | 37.49, 1.13 |
| | **{-7, 7}** | 36.96, 1.21 | 37.33, 0.96 | 37.20, 1.45 | 38.15, 1.84 | 37.11, 0.90 |
| | **{7, -7}** | 38.23, 0.77 | 38.57, 1.27 | 37.80, 1.96 | 37.75, 1.74 | 37.05, 0.92 |
| | **{7, 7}** | 38.13, 1.05 | 38.41, 0.61 | 37.80, 1.44 | 38.14, 0.87 | 37.17, 0.99 |
| **γ** | | 0.018 | 0.024 | 0.009 | 0.055 | 0.063 |
| **λ** | | 0.017 | 0.008 | 0.016 | 0.039 | 0.008 |

**Supplementary Table 1**. Fitted parameters for the psychometric function obtained from the Method of Constant Stimuli (MOCS) experiment at the four tested locations in five subjects with four different combinations of stimulus size and duration. Lapses (λ) and guesses (γ) were modelled as global parameters for the whole MOCS experiment in each eye.

### Results with an alternative temporal impulse response

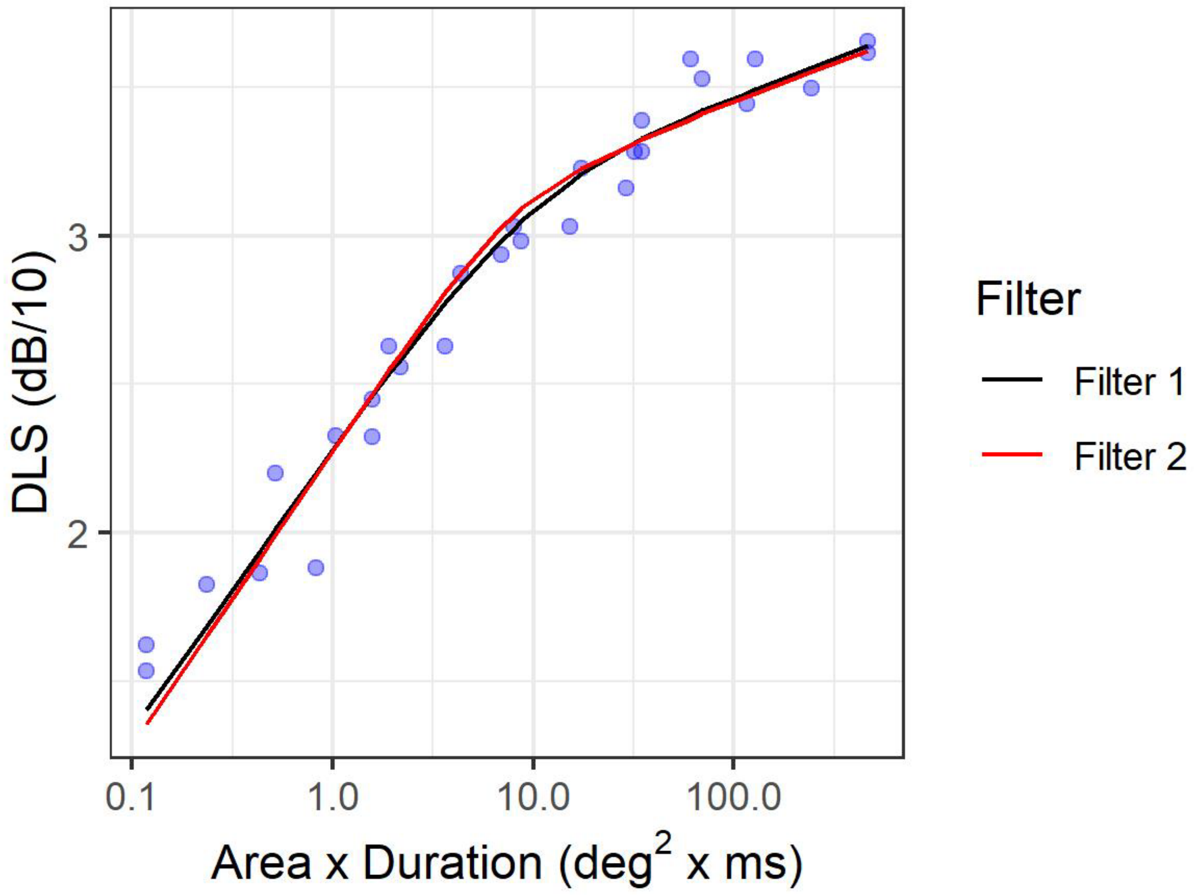


**Figure S3.** Example of the fitted response of the model with the impulse response used in the main calculations (Filter 1) and with the monophasic impulse response used by Gorea and Tyler[1] and described by Watson[2] (Filter 2)
